# Investigation of microglial diversity in a mouse model of Parkinson’s disease pathology

**DOI:** 10.1101/2023.11.23.567809

**Authors:** L Iovino, J VanderZwaag, G Kaur, P Khakpour, V Giusti, A Chiavegato, L Tenorio-Lopes, E Greggio, ME Tremblay, L Civiero

## Abstract

Microglia, the central nervous system resident immune cells, are now recognized to critically impact homeostasis maintenance and contribute to the outcomes of various pathological conditions including Parkinson’s disease (PD). Microglia are heterogenous, with a variety of states recently identified in aging and neurodegenerative disease models, including the ‘disease-associated microglia’ (DAM) which present a selective enrichment of *CLEC7A* encoding the CLEC7A or DECTIN1 protein, and the ‘dark microglia’ (DM) displaying markers of cellular stress at the ultrastructural level. However, the roles of CLEC7A-positive microglia and DM in the pathology of PD have remained largely elusive. By applying immunofluorescence and scanning electron microscopy, we aimed to characterize 1) the CLEC7A -positive cell population, and 2) their possible relationships to DM in a mouse model harboring a G2019S pathogenic mutation of the LRRK2 gene, the most common mutation linked to PD. We examined 18-month-old mice, comparing between LRRK2 G2019S knock-in mice and wild-type controls. In the dorsal striatum, a region affected by PD pathology, extensive ultrastructural features of cellular stress (e.g., endoplasmic reticulum and Golgi apparatus dilation), as well as reduced direct cellular contacts, were observed for microglia from LRRK2 G2019S mice *versus* controls. CLEC7A-positive microglia exhibited extensive phagocytic ultrastructural characteristics in the LRRK2 G2019S mice. Additionally, the LRRK2 G2019S mice presented a higher proportion of DM. Lastly, immunofluorescence and biochemical analysis revealed higher number of CLEC7A-positive cells in Lrrk2 G2019S genotype *versus* controls both in tissues and in primary microglia cells. Of note, CLEC7A-positive cells present a selective enrichment of ameboid morphology and tend to cluster in the pathogenic animal. In summary, we provide novel insights into the involvement of recently-defined microglial states, CLEC7A-positive cells and DM, in the context of LRRK2 G2019S PD pathology.

## Introduction

Microglia form a surveillance network in the central nervous system (CNS), through which they protect the parenchyma and maintain its homeostasis (1). Beside their role in immunity, microglial cells control neuronal proliferation and differentiation, and remodel neuronal circuits throughout the lifetime (2). Only recently, microglia have emerged as a highly heterogenous cellular population (2). Numerous microglial states were identified and classified based on their ultrastructure, morphology, gene and protein expression profile, as well as function, across life stages, CNS regions, and health and disease conditions (2).

Among these microglial states, our group revealed the existence of ‘dark microglia’ (DM) which are distinguished from the other microglia by their markers of cellular stress (e.g., endoplasmic reticulum dilation, mitochondrial cristae alteration) and peculiar electron-density (3,4). DM display a faint and punctiform staining for homeostatic microglial markers, such as IBA,1 P2RY12 and CX3CR1, in contrast to normal or typical microglia showing a strong and diffuse staining for these markers (3). In physiological conditions, DM are almost absent from the young adult mouse brain (3). Conversely, DM are abundant, increasing in number up to 10-fold in mouse models of chronic stress, aging, and neurodegenerative diseases: Alzheimer’s disease (AD) (3), Huntington’s disease (HD) (5), and amyotrophic lateral sclerosis (ALS) (6). Furthermore, DM display a hyper-ramified morphology and their processes mostly localize in close proximity to pre- and post-synaptic elements, suggesting a key role in the remodeling of neuronal circuits, while they express CD11B which is part of the complement receptor 3 and TREM2, both involved in phagocytosis (3). Recently, DM were also identified in post-mortem brains derived from middle-aged and aged human individuals (4), supporting the hypothesis that environmental risk factors such as aging could promote DM appearance (3,4).

Providing key insights into microglial diversity, recent data revealed novel microglial transcriptomic signatures, including the disease associated microglia (DAM) and microglial neurodegenerative phenotype (MGnD), which were identified in different mouse models of neurodegenerative diseases including AD, ALS, and multiple sclerosis (MS) (7,8). Recent transcriptome studies further indicated that the DAM/MGnD express typical microglial markers such as *IBA1* while they upregulate genes such as *Trem2*, *Apo-E*, *Itgax*, *Spp1*, *Lpl* and *Clec7a* (7,8). Interestingly, *Clec7a*-positive microglial clusters were recently identified in aged mice (9–11). *Clec7a* encodes for the protein CLEC7A or DECTIN1, an antifungal receptor involved in both innate and adaptive immunity (12). CLEC7A recognizes the β-(1→3)/(1→6) glucans which are found in the cell wall of several fungi, while its activation has been shown to culminate into a complex and convergent pro-inflammatory response (12). Specifically, CLEC7A activation promotes phagocytosis and the production of reactive oxygen species (ROS) and cytokines such as pro-interleukin 1β (Il-1β), Il-6, and tumor necrosis factor-α (TNF-α) (12). Of note, the role of CLEC7A in the brain and, specifically, in microglia has remained largely elusive, whereas it has been widely studied in peripheral myeloid cells (13–15).

Although in the last years several studies explored microglial states in multiple neurodegenerative diseases (7–11), little is known in Parkinson’s disease (PD) (16). PD is a complex progressive neurodegenerative disease that affects 2–3% of the population ≥ 65 years of age (17). The pathological hallmarks of PD include the premature, selective loss of dopaminergic neurons of the substantia nigra pars compacta (SNpc) projecting to the dorsal striatum (DS), the presence of Lewy Bodies in neuronal cells, and an extended ‘neuroinflammation’ or CNS inflammation (18–23). High levels of TNF-α, IL-1β, ROS, nitric oxide species, and pro-apoptotic proteins have been found in the postmortem SNpc and striatum of PD-affected patients (24,25). Moreover, evidence of astrocyte reactivity has also been observed in the striatum of patients with PD and animal models (24,26). PD is mostly found in sporadic forms, with only 10% of the cases being familial (27). The most common autosomal dominant mutations causing PD are linked to the *LRRK2* gene, which accounts for 4% of the familial cases and 1-2% of the sporadic cases depending on the population and ethnic group (28). The most common *LRRK2* mutation is the pG2019S which causes a two to three-fold increase in the protein kinase activity (28). Recent evidence suggests a possible crosstalk between the LRRK2 and CLEC7A proteins in a transgenic mouse model overexpressing LRRK2 (29). Specifically, *Lrrk2* overexpressing mice displayed a pronounced peripheral inflammatory state caused by an enhanced LRRK2 kinase activity and CLEC7A activation (29). Intriguingly, in this model, the pharmacological inhibition of LRRK2 kinase activity was also found to ameliorate CLEC7A-induced inflammation (29).

To provide novel insights into the pathogenic mechanisms underlying LRRK2-associated PD forms, we characterized microglial diversity, examining possible changes in typical microglia, DM and disease-associated CLEC7A-immunopositive microglial states, within the DS of aged mice harboring the Lrrk2 G2019S pathogenic mutation. Using nanoscale-resolution scanning electron microscopy (ScEM) and ultrastructural analysis, we found that the microglial population from Lrrk2 G2019S knock-in mice exhibits increased cellular stress and reduced number of contacts with the surrounding striatal structures including pre-synaptic axon terminals. As an additional feature of pathology, we detected an increased fraction of DM and the presence of CLEC7A-positive microglia in Lrrk2 G2019S model. *In situ* immunofluorescence (IF) showed that CLEC7A-positive microglia are morphologically different compared to IBA1-positive cells not expressing CLEC7A. In addition, we also demonstrated that IBA1-positive primary microglia derived from Lrrk2 G2019S mice overexpress CLEC7A and that CLEC7A-positive cells display an increased tendency to cluster, similarly to DM, in Lrrk2 G2019S mouse striatum. Together, these data support the coexistence of multiple disease-linked microglial states in the Lrrk2 G2019S mouse striatum, suggesting that microglial phenotypic transformation can contribute to PD pathology.

## Methods

### Animals

C57Bl/6J LRRK2 wild-type (WT) and LRRK2 G2019S knock-in homozygous male and female mice were used. LRRK2 G2019S knock-in mice were obtained from Prof. Michele Morari and the Novartis Institutes for BioMedical Research, Novartis Pharma AG (Basel, Switzerland) (30). Housing and handling of mice were done in compliance with national guidelines. All procedures performed with mice were approved by the Ethical Committee of the University of Padova and the Italian Ministry of Health (license 200/2019).

### Animal Perfusion

18-month-old LRRK2 WT and G2019S mice were anesthetized with xylazine (Rompun®) and Alfaxalone (Alfaxan®). For ScEM and IF experiments, the mice were transcardially perfused with phosphate-buffered saline (PBS (1x solution) 8% NaCl, 0.2% KCl, 1.44% Na_2_HPO_4_, 0.24% KH_2_PO_4_, dissolved in deionized water; pH = 7.4) Afterward, for the ScEM experiments, mice from the same strain were transcardially perfused with ice-cold fixative composed of 4% paraformaldehyde (PFA) and 0.4% glutaraldehyde (both dissolved in PBS, pH = 7.4). Then, the brains were post-fixed for 2 h in 4% PFA. Instead, for the IF experiments, the mice were transcardially perfused with a 4 % PFA (dissolved in PBS, pH 7.4) and the brains were post-fixed in 4 % PFA at 4 °C for 18 h, then they were transferred to two different sucrose solutions (20 % and 30 % dissolved in PBS) at 4 °C for 18 h each. Afterward, coronal brain sections (50 μm thickness for ScEM – 30 μm thickness for IF) were generated using a vibratome (Leica Vibratome VT1000S.) and stored at −20°C in cryoprotectant solution (30% glycerol, 30% ethylene glycol, 40% PBS) until use.

### Immunohistochemistry on Brain Slices for Scanning Electron Microscopy

One cohort of brain sections containing the DS (periventricular, DSp; Supp. Fig. 1) (Bregma 0.02 to 0.86 mm (31)) of Lrrk2 WT and Lrrk2 G2019S C57BL/6J mice (*n*[=[4 per group) were processed for immunostaining against CLEC7A prior to ScEM processing. The chosen sections were quenched with 0.3% H_2_O_2_ (Fisher Scientific, Ottawa, lot# 202762) in PBS for 5 min. Afterwards, the sections were incubated in 0.1% NaBH_4_ in PBS for 30 min followed by 3 washes of 10 min in PBS. The sections were then incubated in a blocking buffer solution containing 10% fetal bovine serum (FBS; Jackson ImmunoResearch Labs, Baltimore, USA cat# 005-000-121), 3% bovine serum albumin (Sigma-Aldrich, Oakville, cat# 9048-46-8,), and 0.05% Triton X-100 in PBS for 1 h at room temperature (RT). They were incubated overnight in blocking buffer solution with the primary Anti-mDectin-1-IgG antibody (1:50; cat# mabg-mdect, Invivogen) at 4°C. The following day, the brain sections were washed with Tris-buffered saline (TBS; 50 mM, pH 7.4) and incubated with a biotinylated goat anti-rat polyclonal secondary antibody (1:300; Jackson ImmunoResearch, Baltimore, USA, cat# 112-065-167) in TBS containing 0.05% Triton X-100 for 90 min at RT. Afterwards, sections were washed in TBS and incubated for 1 h at RT in an avidin-biotin complex solution (ABC; 1:100; Vector Laboratories, Newark, USA, cat# PK-6100) in TBS. The staining was revealed in 0.05% 3,3’-diaminobenzidine (Millipore Sigma, Oakville, USA, cat# D5905-50TAB) activated with 0.015% H_2_O_2_ diluted in Tris buffer (0.05 M, pH 8.0).

### Sample Preparation and Imaging by Scanning Electron Microscopy

The immunostained brain sections containing the DSp (Bregma 0.02 to 0.86 mm (31)) of Lrrk2 WT and Lrrk2 G2019S C57BL/6J mice (*n*[=[1 per group) were processed for ScEM imaging. A second cohort (*n* = 4 per group) of sections was processed for ScEM without immunostaining to prevent interfering with mean gray value analysis and thus facilitate the analysis of DM. The protocol used for the sample preparation for ScEM was recently detailed in St-Pierre et al. (2019). Briefly, the brain sections were rinsed in phosphate buffer (PB) and incubated 1 h in a solution containing equal volumes of 3% potassium ferrocyanide (Sigma-Aldrich, Ontario, Canada, cat# P9387) and 4% osmium tetroxide (EMS, Pensylvannia, USA, cat# 19190) in PB. After washing in PB, the sections were incubated for 20 min in a filtered (0.45μm filter) 1% thiocarbohydrazide solution (diluted in MilliQ water; Sigma-Aldrich, Ontario, Canada, cat# 223220) followed by a second 30 min incubation with 2% aqueous osmium tetroxide (diluted in MilliQ water). The sections were then dehydrated in ascending concentrations of ethanol for 5 min each (2[×[35%, 50%, 70%, 80%, 90%, 3[×[100%) and washed 3 times for 5 min with propylene oxide (Sigma-Aldrich, #cat 110205-18L-C). Next, the sections were embedded in Durcupan resin (20 g component A, 20 g component B, 0.6 g component C, 0.4 g component D; Sigma Canada, Toronto, cat# 44610) overnight. The following day, the resin-infiltrated sections were flat-embedded onto fluoropolymer films (ACLAR^®^, Pennsylvania, USA, Electron Microscopy Sciences, cat# 50425-25) covered with resin and kept at 55 °C in a convection oven for 3 days. After resin was polymerized, areas containing the periventricular striatum were excised and glued onto resin blocks for ultramicrotomy sectioning. Using a Leica ARTOS 3D ultramicrotome, 70-nm thick sections from these areas were generated (4-6 levels,[∼[5–10 µm apart) and collected on silicon wafers (EMS, Pennsylvania) for imaging on a Zeiss Crossbeam 350 focused ion beam-scanning electron microscope (FIB-SEM). Using the software Zeiss Atlas 5 (Fibics, Ottawa), images of microglia were randomly acquired in the DSp at a resolution of 5 nm per pixel and exported in .tif for ultrastructural analysis.

### Quantitative Ultrastructural Analysis of Microglia

For the ultrastructural analysis, 10–19 microglia per animal (*n*[=[4 per group) imaged in the DSp at a resolution of 5 nm per pixel were examined. The images were blinded to the experimental condition prior to analysis to prevent bias. To quantify ultrastructural changes, we analyzed 102 microglial cell bodies in total (48–52 per experimental condition), a sample size considered sufficient to obtain statistical power based on the G*Power software V3.1 (effect size of 0.75 and power of 0.95 estimated to 100 individual cells). A similar effect size was previously used to quantitatively assess microglial ultrastructure in mice (32). The identification of microglia and their cytoplasmic content was previously described in detail (33). Briefly, microglia were differentiated from neurons and astrocytes by their size and nuclear heterochromatin pattern, and further distinguished from oligodendrocytes by their long and narrow stretches of endoplasmic reticulum, diverse inclusions (e.g., lysosomes) dispersed heterogeneously in their cytoplasm, and association with pockets of extracellular space, among other key distinctive features (33). The identifications of microglia were agreed upon by three trained, separate, and blinded observers to provide consensus and prevent bias. Examples images of microglia in both LRRK2 WT and G2019S mice are provided (Supp. Fig. 2).

Microglia were considered to be in contact with other parenchymal structures such as the vasculature or other surrounding cell bodies (e.g., astrocytes, neurons, oligodendrocytes) if the distance between their cell body plasma membrane and associated structure’s membrane (e.g., synapse, neuronal cell body) was under 15 nm. Microglial contacts with synaptic elements were categorized as axon terminals, post-synaptic dendritic spines or synapses (when the two elements and synaptic cleft were contacted simultaneously (4)). Axon terminals were defined by their > 5 synaptic vesicles, while post-synaptic dendrites were recognized by their postsynaptic density (34,35). Myelinated axons were identified by their electron-dense membrane sheaths (36). Extracellular digestion and extracellular space consisted of electron-lucent space pockets devoid of a membrane delineation located next to the microglia and containing (degeneration) or not (space) partially-digested cellular elements (37). One-to-five micron diameter granular aggregates were found throughout the striatum of mice and were hypothesized to be protein deposition resembling previous ultrastructural observations of alpha-synuclein (38).

Lysosomes were identified by their circular and homogenous (primary) or heterogenous (secondary, tertiary) contents. Secondary lysosomes were larger than primary lysosomes and displayed a more heterogenous appearance. Tertiary lysosomes were larger than primary and secondary lysosomes, and primarily displayed a heterogeneous appearance, often possessing large lipid bodies in addition to phagosomes (39,40). Endoplasmic reticulum (ER) cisternae were characterized by their long and narrow stretches found inside the microglial cytoplasm (33). Dilated ER and Golgi bodies were identified by their swollen electron-lucent appearance, with a cisternae width over 100 nm indicating dilation (4). Altered ER and Golgi bodies were characterized by deterioration or disruption of membranes, typically with other cellular materials contained in the cisternae. The term “dystrophic” was used to describe the subtotal of all ER and Golgi not identified as healthy. Mitochondria were identified by their electron-dense appearance, double membrane, numerous cristae, and round shape. Altered mitochondria were characterized by the deterioration of the outer membrane, degradation of the cristae (electron-lucent pockets), or a “holy” shape, which refers to mitochondria forming a donut structure (41). Mitochondria with a length over 1000 nm were defined as elongated (42). The term “dystrophic” was used to describe the subtotal of all mitochondria not identified as healthy.

To evaluate the microglial area, perimeter and morphology, each microglial cell body was traced using the polygon tool in ImageJ, and various shape descriptors such as aspect ratio, area, perimeter, solidity, and circularity were analyzed as previously described (36,39,43,44). Aspect ratio refers to the ratio of the height to the width of the microglial cell body. Solidity is a measure of irregular shapes, calculated by dividing the cell area by the convex area. The closer the solidity value is to 1, the less irregular the microglial cell body shape is. Circularity is calculated as 4π times the area over the perimeter squared and is used to assess the cell shape where a value closer to 1 indicates a rounder shape and a value near 0 indicates an elongated shape. To evaluate lysosomal area, each mature (secondary or tertiary) lysosome per cell was traced using the polygon tool in Image J and area was analyzed. The percentage of soma area containing mature lysosomes was calculated using the following:

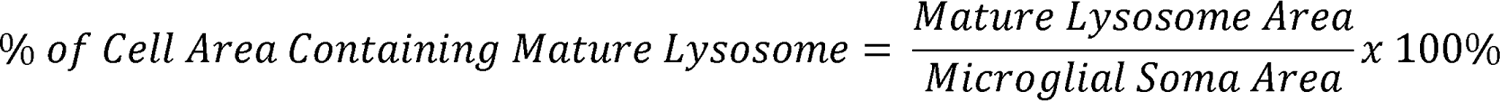

To evaluate the percentage of the perimeter of each microglial cell body contacting extracellular degeneration or space pockets, the length of each section contacting electron-lucent space devoid of a membrane delineation located next to the microglia and containing (degeneration) or not (space) partially digested cellular elements was measured using the polygon tool in Image J. The sums of all lengths of sections matching this description were divided by total cell perimeter in the following:

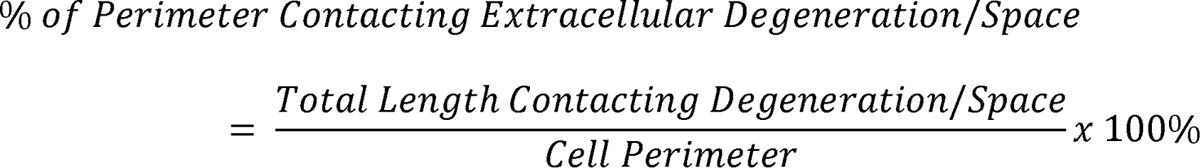

### Quantitative Analysis of Microglial Integrated Density Ratio

As previous research has identified DM (and intermediate microglia, presenting features of both typical microglia and DM) as an abundant cell population in aging and neurodegenerative disease pathology (3,39,45,46), a quantitative approach to measuring the shade, or “darkness”, of microglia on electron micrographs was next employed. Using ImageJ, the integrated density of microglial cytoplasm and integrated density of nearby astrocytic cytoplasm (in either cell body or process) was measured. The relative “darkness” of each microglial cell body was thus calculated using the following:

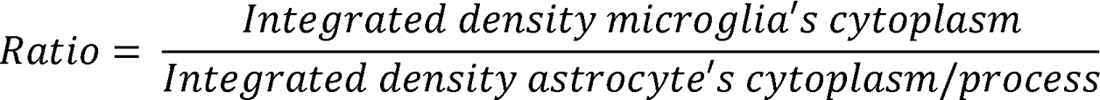

Then integrated dentistry ratio data were further transformed by categorization into the following ranges > 0.875, 0.875-0.750, and < 0.750 arbitrary units (a.u.) for group-wise comparison. These ranges were chosen to match previous qualitative categorization of typical, intermediate, and dark microglia, respectively (3,39,45,46). In short, intermediate microglia can be distinguished from typical microglia based on their more electron-dense cytoplasm and heterochromatin pattern, accompanied by intermediate markers of cellular stress such as ER dilation (3,39,45,46). DM can be further distinguished from typical and intermediate microglia based on their cytoplasm and nucleus, which appear even more electron-dense, and frequent absence of typical nuclear heterochromatin pattern, in addition to their various markers of cellular stress including ER dilation and mitochondrial alterations (3,4,39,41,45). Proportion of each cell category (typical, intermediate, and dark) in each treatment group was calculated using the following formula:

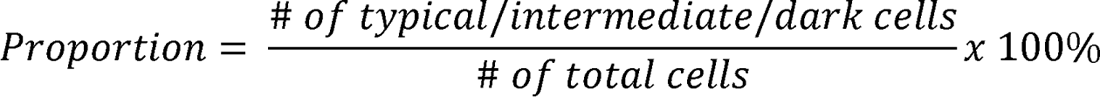

### Primary Microglial Cells

Primary microglial cells derived from Lrrk2 WT and G2019S postnatal day (PND) 0–4 sex-matched mouse pups were prepared as previously described (47) with a few modifications. Briefly, tissue was mechanically dissociated in cold PBS (Biowest, #L0615) and the meninges were removed. Cerebral cortex was selected as a region of interest to reach the required microglial cell yield. Then, the tissues were transferred to 15 mL Falcon tubes containing 1 mL of Dulbecco’s Modified Eagle Medium (DMEM, Biowest, #L0103) supplemented with 10% FBS (Corning) and 100 U/mL Penicillin + 100 µg/mL Streptomycin (Pen-Strep; Life Technologies). The tissues were mechanically dissociated by pipetting 25 times. Afterward, 11 mL of supplemented DMEM were added and the cellular suspension was allowed to settle for 5 min. The top fraction (10 mL) was collected and centrifuged for 10 min at 200 g. The pellets were re-suspended in supplemented medium and plated on poly-L-lysine (0.1 mg/ml, Sigma Aldrich)-coated T-75 flask. After 24 h, flasks were washed three times with PBS and the medium was replaced with fresh medium containing complete DMEM supplemented with 10% L929-conditioned medium. After 12 days, cells were shacked off and used for the Western blot or IF experiments.

### Immunofluorescence on Primary Microglial Cells and Data Analysis

Primary microglial cells were seeded at a density of 2 x 10^5^ cells on 12 mm glass coverslips (VWR) coated with poly-L-lysine (0.1 mg/mL). After 24 h, they were fixed in 4 % PFA for 20 min at RT. Thereafter, the cells were washed in PBS and permeabilized for 20 min in PBS containing 0.1 % Triton X-100. Then, primary microglial cells were incubated for 1 h at RT in a blocking solution containing 5 % FBS in PBS and incubated overnight with a mixture of the following primary antibodies diluted in the blocking solution: rabbit anti-IBA1 (1:300 – Wako; 019-19741) and rat anti-mDectin1 (1:50) antibodies. The following day, microglial cells were incubated for 1 h at RT with a mixture of anti-rabbit Alexa Fluor 488 (Invitrogen; #A-11008) and anti-rat Alexa Fluor 647 (Invitrogen; # A-21247) antibodies diluted 1:200 in the blocking solution. Nuclei were counterstained for 5 min with Hoechst (1:10000 in PBS; Invitrogen), and after washing, coverslips were mounted using Mowiol (Calbiochem).

Stained primary microglial cells (from *n* = 5 independent cultures) were imaged at 8-bit intensity resolution over 1024 x 1024 pixels on a Leica SP5 confocal microscope using a HC PL FLUOTAR 40x/0.70 dry objective. We counted the number of CLEC7A^+^ (IBA1^+^ and IBA1^-^), IBA1^+^ (CLEC7A^+^ and CLEC7A^-^), and double-positive (IBA1^+^/CLEC7A^+^) primary microglial cells, and we normalized the number of counted cells to the total number of nuclei in the field. Following the same method, we also quantified the number of CLEC7A^+^/IBA1^-^, as well as the proportion of IBA1^+^/CLEC7A^-^ primary microglial cells. Finally, the correlation between the intensity of IBA1 and CLEC7A was evaluated using the JACoP plugin in ImageJ. Briefly, for each image, the Pearson’s correlation r between IBA1 and CLEC7A was determined as previously described (48).

### Immunofluorescence on Brain Slices and Data Analysis

Sections containing the DS (Bregma 1.2 to 0.5 mm (31)) were selected and were rinsed in PBS. Sections were quenched for 5 min with a 0.3% H_2_O_2_ (Carl Roth Gmbh & Co. Kg) in PBS. After three washes with PBS, the sections were treated with 0.1% NaBH_4_ (Biochemika, #71997) in PBS for 15 min at RT to quench intrinsic autofluorescence. Sections were then washed in PBS followed by permeabilization and saturation for 1 h at RT in a blocking solution containing 2 % BSA, 15% goat serum, 0.25 % gelatin, 0.2 % glycine and 0.5 % Triton-X100 in PBS. Sections were next incubated overnight with primary antibodies diluted in blocking solution. The following primary antibodies were used: rabbit anti-IBA1 (1:300) and rat anti-mDectin1 (1:50). The following day, sections were washed and incubated for 1 h at RT with the appropriate secondary antibodies diluted 1:200 in blocking solution: anti-rabbit Alexa Fluor 488 and anti-rat Alexa Fluor 647 fluorophores. Upon incubation, the sections were washed with PBS, counterstained with Hoechst (1:10000 in PBS) to label nuclei and mounted using Mowiol (ThermoFisher) on glass microscopy slides.

For each animal (*n* = 4 per genotype), two brain sections were used and for each section, at least 3 images were acquired. Images were acquired at 8-bit intensity resolution over 1024 x 1024 pixels on a Leica SP5 confocal microscope using a HC PL FLUOTAR 20X/0.50 dry objective. To characterize the overall proportion of IBA1^+^ and CLEC7A^+^ cells, we firstly split the image in different channels. Then, we counted the number of CLEC7A^+^ (IBA1^+^ and IBA1^-^), IBA1^+^ (CLEC7A^+^ and CLEC7A^-^), and double-positive cells (IBA1^+^/CLEC7A^+^) and we normalized them to the total number of counted cells. We also quantified the proportion of CLEC7A^+^/IBA1^-^ cells and the proportion of IBA1^+^/CLEC7A^-^ cells. The number of total counted cells used for the normalization were estimated as:

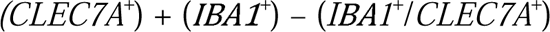

To determine the number of clusters comprising at least three CLEC7A^+^ cells (IBA1^+^ and IBA1^-^), we acquired at least eight images per condition at 8-bit intensity resolution over 1024 x 1024 pixels on a Leica SP5 confocal microscope using a HC PL FLUOTAR 40x/0.70 dry objective. Finally, for the morphological analysis, 16 IBA1^+^/CLEC7A^-^ and 16 IBA1^+^/CLEC7A^+^ cells were analyzed. Images were acquired at 8-bit intensity resolution over 1024 x 1024 pixels on a Leica SP5 confocal microscope using a HC PL FLUOTAR 63X/0.80 oil objective. Only IBA1^+^/CLEC7A^-^ as well as double-positive cells (IBA1^+^/CLEC7A^+^) completely captured within the z-stacks were considered for the analysis. The arborization was outlined using the polygon tool of the ImageJ software and the area, perimeter, circularity, and solidity were measured as previously described (49,50). Moreover, the Sholl analysis was performed (step size = 1 μm; radius = 100 μm), as described in (51), by using the tracing on the microglial cell performed with the NeuronJ plugin (version 1.4.3). Finally, for each cell, the integrated density of IBA1 and CLEC7A was measured. Briefly, using the same regions of interest (ROIs) used for the morphological analysis, the integrated density for both channels was calculated blind to the experimental conditions using ImageJ software.

### Immunoblotting

Immunoblotting was performed on both primary microglial cells (from *n* = 6 independent cultures) and cortex/striatum tissues derived from 18-month-old Lrrk2 WT and Lrrk2 G2019S male and female mice (*n* = 3 per genotype). Briefly, upon cervical dislocation, the brain was retrieved from the adult mice. Cortices and striatum were then collected by dissection under a stereomicroscope and lysed using tissue grinders (26).

Dissected primary cortical microglial cells or murine cortex/striata derived from LRRK2 WT and LRRK2 G2019S knock-in mice were lysed in radioimmunoprecipitation assay buffer (RIPA; 20 mM Tris-HCl pH 7.5, 150 mM NaCl, 1 mM Na_2_EDTA, 1 mM EGTA, 1% NP-40, 1% sodium deoxycholate, 2.5 mM 30 sodium pyrophosphate, 1 mM β-glycerophosphate, 1 mM Na_3_VO_4_, deionized water) containing 1 % protease inhibitor cocktail (Sigma-Aldrich). Protein concentration was measured using the Pierce® BCA Protein Assay Kit following the manufacturer’s instructions (Thermo Scientific). The samples were resolved by electrophoresis on ExpressPlus PAGE precast gels 4–20% (GeneScript), according to the manufacturer’s instructions. After electrophoresis, protein samples were transferred to polyvinylidene fluoride (PVDF) membranes (Bio-Rad) through a Trans-Blot TurboTM Transfer System (Bio-Rad) in semi-dry conditions, with the 1× transfer buffer (Bio-Rad) at 25 V for 20 min. Membranes were incubated in Tris-buffered saline containing 0.1 % Tween (TBS-T) and 5 % skimmed milk for 1 h at RT, then incubated overnight with primary antibodies diluted in TBS-T plus 5 % skimmed milk. The following primary antibodies were used: rabbit anti-IBA1 (1:1000), mouse anti-HSP70 (1:5000 – Sigma Aldrich; H5147), rabbit anti-CLEC7A (1:500 – Invitrogen; PA5-97284). Membranes were subsequently rinsed and incubated for 1 h at RT with the appropriate horseradish-peroxidase (HRP)-conjugated secondary antibodies (Invitrogen). The signal was visualized using Immobilon® Forte Western HRP Substrate (Millipore) and the VWR® Imager Chemi Premium. Images were acquired and processed with ImageJ software to quantify total intensity of each single band by performing a densitometry analysis. The intensity of the band of the protein of interest was then normalized to the intensity of the corresponding housekeeping protein.

### Image Preparation and Statistical Analyses

All the images were prepared for illustration using Adobe Illustrator CS6. Statistical analyses were performed in Prism 7 (GraphPad). Data are expressed as mean ± standard error of the mean (SEM). Gaussian distribution was assessed by D’Agostino & Pearson omnibus and Shapiro-Wilk normality tests. The ROUT method (Q = 1 %) was applied before statistical analysis to remove outliers on data following a Gaussian distribution. Statistical analysis on data including two independent groups was performed with two-tailed unpaired Student’s t-tests (Gaussian distribution) or Mann-Whitney tests (non-Gaussian distribution). To compare the effects of different variables on two different groups, we used two-way ANOVAs followed by Tukey’s multiple comparisons tests. ANOVA p values are reported in the main text. Levels of significance are defined as * p ≤ 0.05; ** p ≤ 0.01; *** p ≤ 0.001; **** p ≤ 0.0001.

Specific to the quantitative analysis of microglial integrated density ratio, the raw integrated density ratio data were checked for Gaussian distribution and then analyzed for group-wise differences using a Mann-Whitney test. To compare differences in the prevalence of each ultrastructural microglial state (typical, intermediate and dark microglia) between the two analyzed groups (Lrrk2 WT and G2019S), each image within the SEM dataset was categorized based on its integrated density and data was dummy coded with 1 as the in-group (e.g., dark) and 0 as the out-group (e.g., intermediate and typical). Then a Mann-Whitney U test was performed to compare differences in the abundance of microglia that were typical (> 0.875 a.u.), intermediate (0.875–0.750 a.u.), and dark (< 0.750 a.u.) in the DSp between Lrrk2 WT and G2019S gene knock-in mice. Levels of significance are reported consistently throughout manuscript.

## Results

### Lrrk2 G2019S Microglia Display Extensive Ultrastructural Features of Stress, Decreases in Direct Contacts with Axon Terminals and Synapses, and Increased Contacts with Extracellular Digestion and Space

To provide insights into microglial involvement in LRRK2-linked PD pathology, microglial cell bodies in the DSp of both 18-month-old Lrrk2 WT and Lrrk2 G2019S mice (Suppl. Fig. 1) were first examined for their shape descriptors by ScEM. We did not find any differences in aspect ratio, area, perimeter, solidity and circulatory (Supp. Table 1). We next examined ultrastructural markers of cellular stress (e.g., ER and Golgi dilation, elongated mitochondria) by ScEM (Fig. 1). The Lrrk2 WT microglia displayed significantly greater number of total ER (U = 920.5; p = 0.0105) and healthy ER (U = 773.5; p = 0.0003) compared to Lrrk2 G2019S microglia (Fig. 1D-E). By contrast, Lrrk2 G2019S microglia displayed a significantly greater number of dilated ER (U = 807.5; p = 0.0005) and a significantly higher percentage of total ER characterized as dystrophic (U = 818.5; p = 0.0007) when compared to Lrrk2 WT microglia (Fig. 1F-G). Similarly, Lrrk2 G2019S microglia presented insignificant changes in total Golgi bodies (U = 1204; p = 0.4458), but significantly fewer healthy Golgi bodies (U = 916.0; p < 0.0001) compared to Lrrk2 WT microglia (Fig. 1H-I). Lrrk2 G2019S microglia had significantly greater number of dilated Golgi bodies (U = 1031; p = 0.0032) and a significantly higher percentage of total Golgi bodies displaying signs of dystrophy (U = 1031; p = 0.0032) when compared to Lrrk2 WT microglia (Fig. 1J-K). The total numbers of dilated Golgi bodies and ER in Lrrk2 G2019S microglia were also significantly higher (U = 717.5; p < 0.0001) compared to Lrrk2 WT microglia (Fig. 1L).

**Fig. 1:**
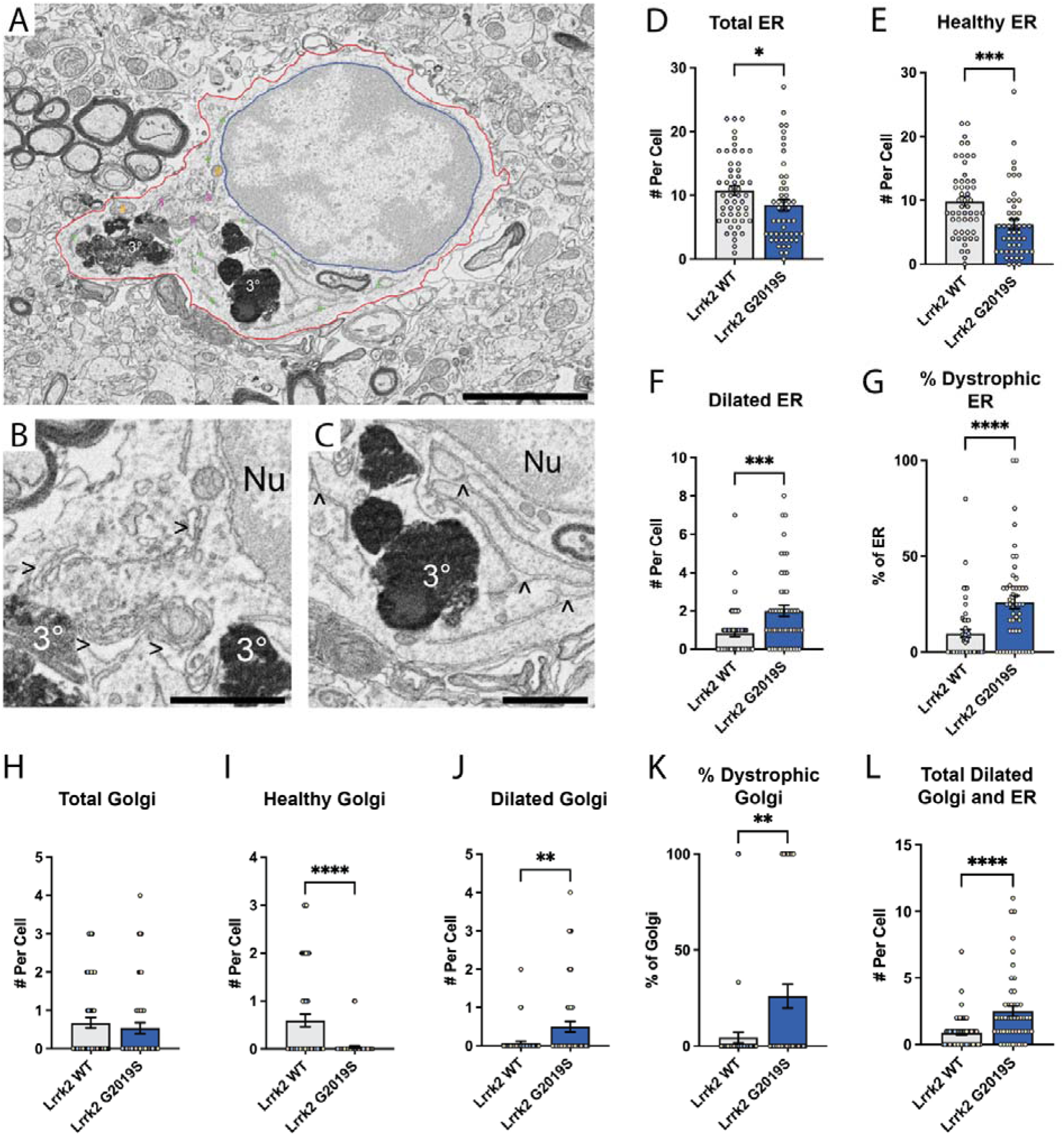
Impact of Lrrk2 G2019S pathological mutation on the structural characteristics of the endoplasmic reticulum (ER) and Golgi apparati in microglial cell bodies from the DSp. (A) Example of parenchymal Lrrk2 G2019S microglial cell with classic microglial structural characteristics: tertiary lysosomes, extensive ER, and nuclear heterochromatin pattern [scale bar = 3 μm]. (B-C) Magnified image of dilated Golgi bodies (>) and dilated ER (^) [scale bar = 1 μm]. (D-F) Quantification of total ER, healthy ER, and dilated ER identified per cell analyzed in Lrrk2 WT and G2019S mice. (G) Calculation of percentage of total ER that were identified as dystrophic (subtotal of dilated ER and altered ER) per cell analyzed. (H-J) Quantification of total Golgi, healthy Golgi, and dilated Golgi identified per cell analyzed in Lrrk2 WT and G2019S mice. (K) Calculation of percentage of total Golgi that were described as dystrophic (subtotal of dilated Golgi and altered Golgi) per cell analyzed. (L) Quantification of total dilated ER and dilated Golgi identified per cell analyzed in Lrrk2 WT and G2019S mice. Quantitative data are shown as individual dots with mean[±[S.E.M. * p[<[0.05, ** p[<[0.01, *** p < 0.001, **** p < 0.0001 using a non-parametric Mann–Whitney test. Statistical tests were performed on *n*LJ=[10-19 microglia per animal with N[=[4 mice/group, for a total of 102 microglial cell bodies analyzed. Red outline[=[plasma membrane, blue outline[=[nuclear membrane, orange octothorpe[=[mitochondria, green plus sign[=[ER, purple ampersand[= Golgi apparatus, > = dilated Golgi apparati, ^ = dilated ER, 3°[=[tertiary lysosome, Nu = nucleus.

Subsequently, microglia in the DSp of both Lrrk2 G2019S and WT mice were examined at the ultrastructural level to quantify direct contacts with surrounding parenchymal structures (e.g., neuronal cell bodies, synapses, myelinated axons). No significant decreases in the number of contacts with neuronal cell bodies (U = 1159; p = 0.1880) were found for Lrrk2 G2019S microglia compared to Lrrk2 WT microglia (Fig. 2C). However, contacts with pre-synaptic axon terminals (U = 664.0; p < 0.0001) and synapses (U = 1050; p = 0.0104) were significantly decreased in Lrrk2 G2019S microglia compared to Lrrk2 WT microglia (Fig. 2D-E). A similar, yet non-significant, tendency toward reduced contacts with post-synaptic dendritic spines (U = 1252; p = 0.6783) was found in Lrrk2 G2019S microglia (Fig. 2F). Consequently, Lrrk2 G2019S microglia had significantly fewer direct contacts with all synaptic structures (U = 566.0; p < 0.0001) when compared to Lrrk2 WT microglia (Fig. 2G). Overall, Lrrk2 G2019S microglia presented significantly decreased total cellular contacts with surrounding parenchymal structures (U = 708.5; p < 0.0001) compared to Lrrk2 WT microglia (Fig. 2H). Lrrk2 G2019S microglia also displayed an increased percentage of their total perimeter surrounding extracellular space pockets containing debris (U = 187.5; p < 0.0001) compared to Lrrk2 WT microglia (Fig. 2I). Interestingly, Lrrk2 G2019S *versus* Lrrk2 WT microglia further showed a significant increase in their contacts with myelinated axons (U = 1003; p = 0.0207) (Fig. 2J).

**Fig. 2:**
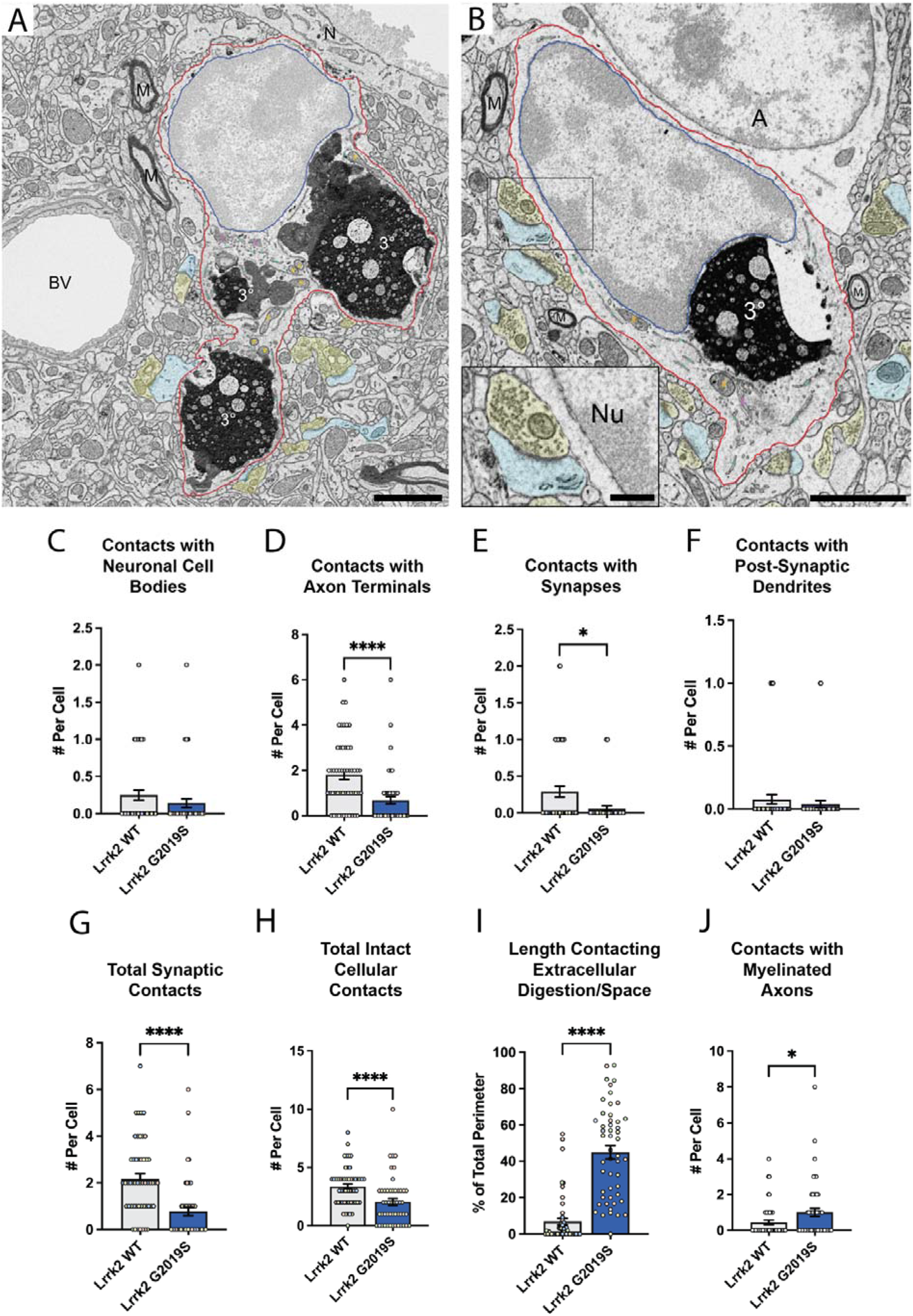
Impact of Lrrk2 G2019S mutation on the direct contacts with neural structures for microglial cell bodies from the DSp. (A) Example of Lrrk2 WT parenchymal microglia in the periventricular striatum presenting typical microglial structural characteristics: tertiary lysosomes, extensive endoplasmic reticulum (ER), and nuclear heterochromatin pattern [scale bar = 2μm]. (B) Secondary example of Lrrk2 WT microglia in the periventricular striatum situated in a satellite position on an astrocyte, with a magnified image showing an example of direct contact with an excitatory synapse [scale bar = 2 μm; inset scale bar = 0.5 μm]. (C-F) Quantification of microglial contacts with neuronal cell bodies, axon terminals, synapses, and post-synaptic dendrites identified per cell analyzed in Lrrk2 WT and G2019S mice. (G) Quantification of microglial contacts with all synaptic structures (axon terminals, synapses, and post-synaptic dendrites) identified per cell analyzed in Lrrk2 WT and G2019S mice. (H) Quantification of total microglial intact contacts with all parenchymal structures (synaptic structures, other cell types, blood vessels, myelin, etc.) identified per cell analyzed in Lrrk2 WT and G2019S mice. (I) Quantification of the percentage of total perimeter of each cell body contacting extracellular digestion/space identified per cell analyzed in Lrrk2 WT and G2019S mice. (J) Quantification of total microglial contacts with all myelinated axons per cell analyzed in Lrrk2 WT and G2019S mice. Quantitative data are shown as individual dots with mean[±[S.E.M. * p[<[0.05, ** p[<[0.01, *** p < 0.001, **** p < 0.0001 using a non-parametric Mann–Whitney test. Statistical tests were performed on *n*[=[10-19 microglia per animal with N[=[4 mice/group, for a total of 102 microglial cell bodies analyzed. Red outline[=[plasma membrane, blue outline[=[nuclear membrane, yellow pseudocoloring = axon terminal, blue pseudocoloring = post-synaptic dendrite, orange octothorpe[=[mitochondria, green plus sign[=[ER, purple ampersand[= Golgi apparatus, 3°[=[tertiary lysosome, A = astrocyte cell body, BV = blood vessel, N = neuronal cell body, Nu = nucleus, M = myelinated axon.

Overall, our data indicate that microglia in the DSp of Lrrk2 G2019S mice displayed significantly more extensive ultrastructural markers of cellular stress associated with a significantly reduced prevalence of direct contacts with surrounding parenchymal structures, particularly axon terminals and synapses.

### Lrrk2 G2019S Mice Exhibit Multiple Disease-linked Microglial States

We next examined DM, a unique state of microglia visualized through electron microscopy that were previously shown to become abundant with aging and neurodegenerative disease pathology (3,4,45). These DM were frequently observed in the DSp of the Lrrk2 G2019S model (Fig. 3A-C). To quantify the abundance of dark and intermediate microglia, which display intermediate features between typical and dark microglia (e.g., cellular stress markers, electron density), microglial cell bodies located in the DSp of Lrrk2 G2019S *versus* WT mice were quantitatively analyzed. The microglial cytoplasmic integrated density was compared to nearby astrocytic cytoplasm integrated density. Overall, microglia in Lrrk2 G2019S mice had lower mean value and broader range of integrated density ratio compared to those in Lrrk2 WT mice; however, this difference was statistically insignificant when examining all microglial cells together (U = 1064, p = 0.201) (Fig. 3D). After the integrated density ratio data was categorized into typical, intermediate, and dark groups, this analysis revealed more typical microglia in Lrrk2 WT (86.54% of cells) mice compared to Lrrk2 G2019S mice (64.58% of cells; U = 974, p = 0.018) (Fig. 3A-C, E). Similarly, a significant increase in the abundance of DM was found in Lrrk2 G2019S (16.67% of cells) mice compared to WT controls (0.000% of cells; U = 1040, p = 0.002) (Fig.3A-C, E). No significant difference in the quantity of intermediate microglia was found between Lrrk2 G2019S (18.75% of cells) and WT (13.46% of cells; (U = 1182, p = 0.588) (Fig. 3E). The microglia classified as dark typically were fully or nearly lacking heterochromatic pattern and displayed signs of stress, particularly dilated Golgi body and mitochondrial cristae alterations (Fig. 3C). These dark microglia made frequent contacts with surrounding parenchymal elements such as myelinated axons (Fig. 3C).

**Fig. 3:**
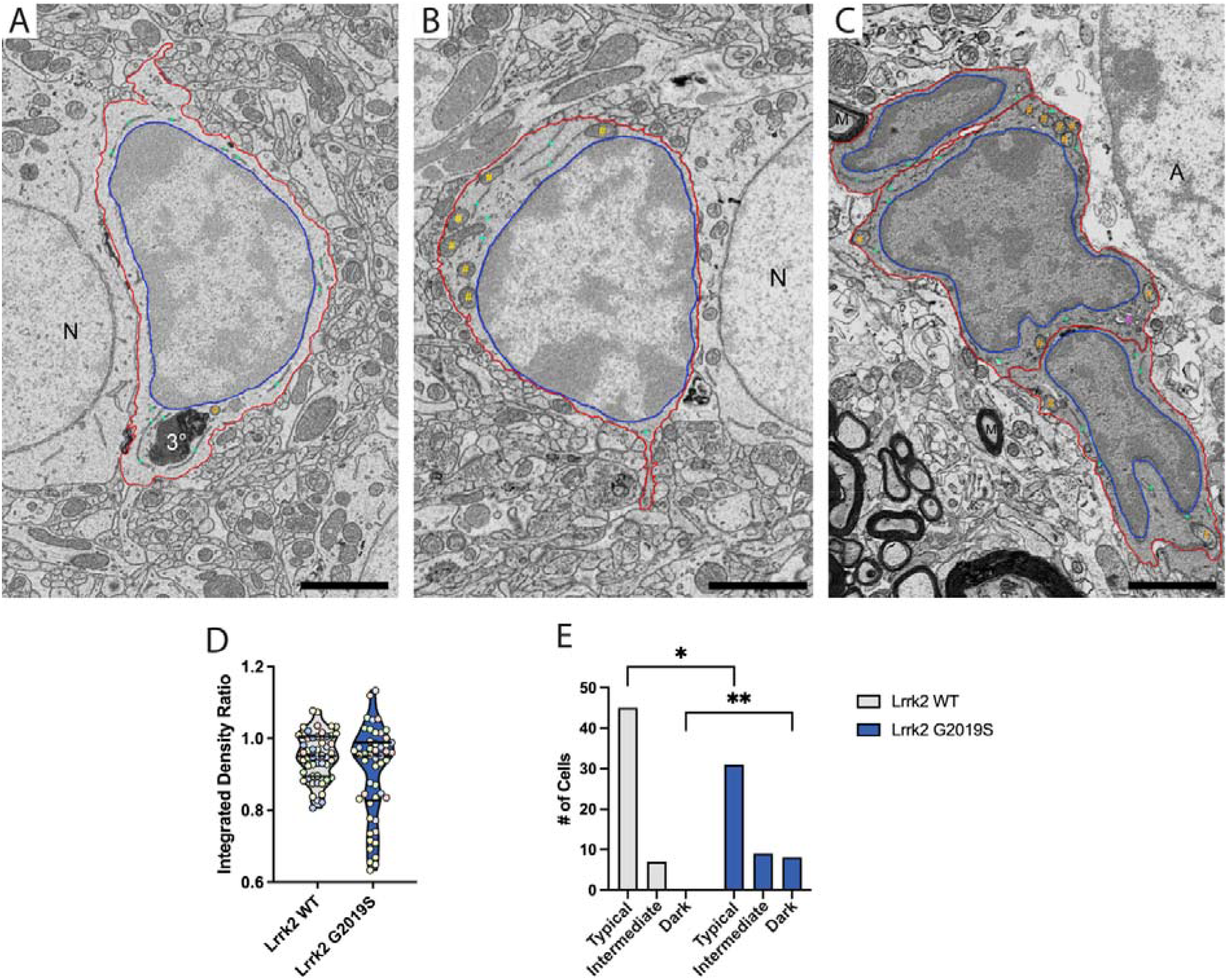
Impact of Lrrk2 G2019S mutation on dark, intermediate and typical microglia identified via their integrated density ratio in the DSp. (A) Example of Lrrk2 WT parenchymal microglia in DSp with a relative difference in integrated density greater than 0.875 a.u. (actual value = 0.970 a.u.) when compared to nearby astrocytic processes and thus categorized as typical [scale bar = 2 μm]. (B) Example of Lrrk2 G2019S microglia in the DSp with a relative difference in integrated density between the values of 0.875 a.u. and 0.750 a.u. (actual value = 0.844 a.u.) when compared to nearby astrocytic processes and thus categorized as intermediate [scale bar = 2 μm]. (C) Example of Lrrk2 G2019S microglia in the DSp with a relative difference in integrated density less than 0.750 a.u. (actual value = 0.656 a.u.) when compared to nearby astrocytic processes and thus categorized as dark [scale bar = 2 μm]. (D) Truncated violin plot of integrated density ratio of microglia located in the DSp of Lrrk2 WT and G2019S mice; group-wise comparison performed using non-parametric, Mann–Whitney test with median displayed as dotted line, quartiles as solid line, and individual data points. (E) Frequency distribution of typical (>0.875 a.u.), intermediate (0.875-0.750 a.u.), and dark (<0.750 a.u.) in DSp of Lrrk2 WT and G2019S n mice. * = p < 0.05, ** = p < 0.001 using a non-parametric Mann-Whitney U test. Statistical tests were performed on *n*[=[10-19 microglia per animal with N[=[4 mice/group, for a total of 102 microglial cell bodies analyzed. Red outline[=[plasma membrane, blue outline[=[nuclear membrane, orange octothorpe[=[mitochondria, green plus sign[=[endoplasmic reticulum, purple ampersand[= Golgi apparatus, 3°[=[tertiary lysosome, A = astrocyte cell body, N = neuronal cell body, Nu = nucleus, M = myelinated axon.

To further investigate CLEC7A^+^ disease associated microglial states in the Lrrk2 G2019S model, sections containing the DSp were immunostained for CLEC7A with peroxidase (which produces a diffusible electron dense precipitate) and imaged on the ScEM. With the stringent immunostaining conditions for electron microscopy, CLEC7A^+^ microglia were found exclusively in Lrrk2 G2019S mice where the immunostaining was broadly filling their cytoplasm and decorating their plasma membrane (Fig. 4A-C). The CLEC7A^+^ cell bodies frequently possessed ultrastructural features of cellular stress including ER and Golgi body dilation (Fig. 4A-C). Additionally, CLEC7A^+^ microglia indicated extensive features of phagocytic activity, including accumulation of lipids and myelin (Fig. 4C, E), cholesterol crystals (Fig. 4A, D), autophagosomes (Fig. 4C), and a tight association with extracellular space and digestion (Fig. 4). Interestingly, CLEC7A^+^ microglia were also seen with large ingested protein aggregates, likely associated with PD and other neurodegenerative disease pathology (Fig. 4B, F; (53)). While we did not observe CLEC7A^+^ cells with a dark cytoplasm, which could have been masked by the peroxidase immunostaining, some displayed features of intermediate microglia (Suppl. Fig. 3).

**Fig. 4:**
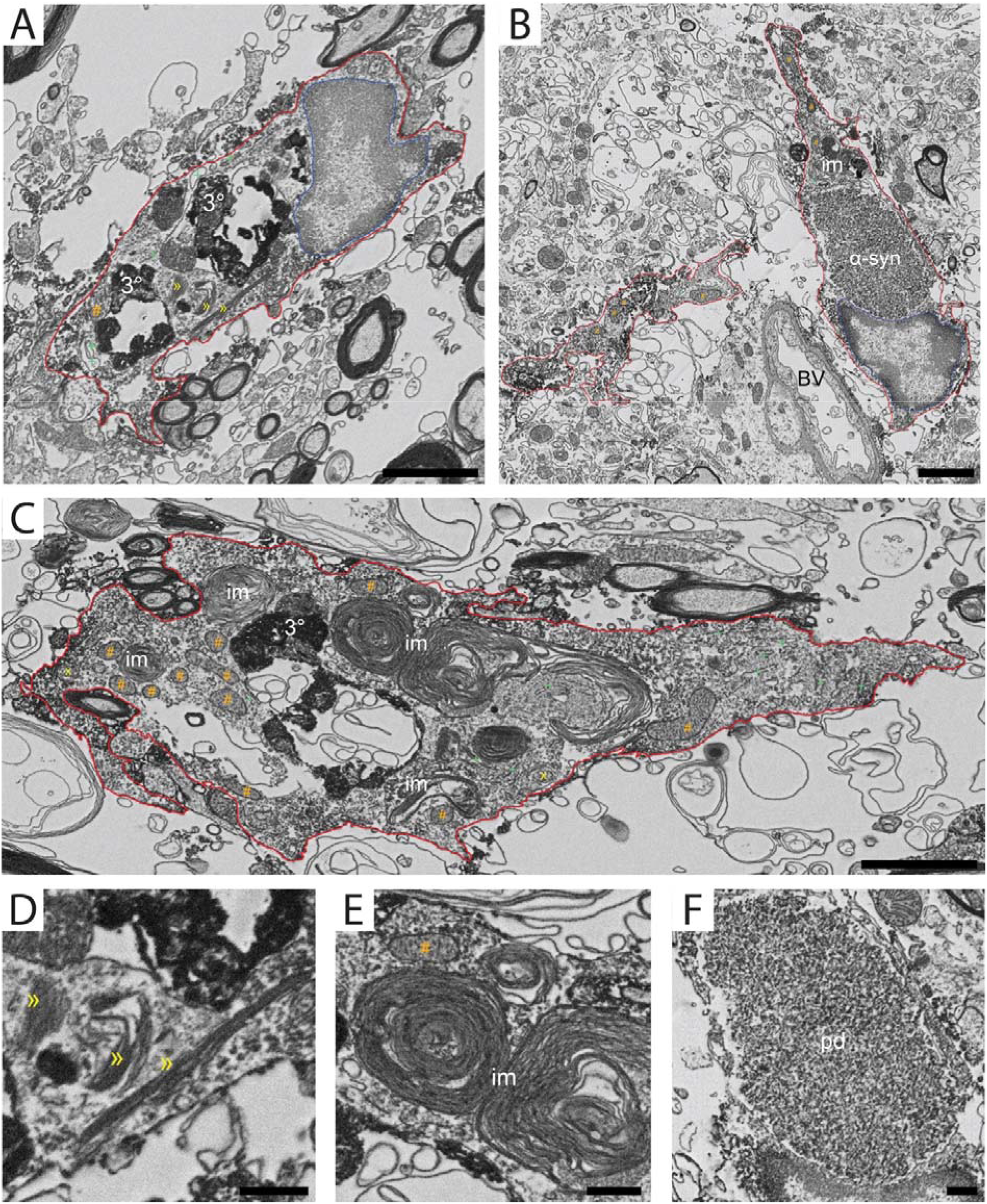
Impact of Lrrk2 G2019S mutation on CLEC7A-immunopositive microglia in the DSp. (A) Example of CLEC7A-immunopositive microglial cell body located in the DSp of a Lrrk2 G2019S mouse which contains cholesterol crystals (») and tertiary lysosomes (3°) [scale bar = 2 μm]. (B) Example of CLEC7A-immunopositive microglial cell body and nearby microglial process with ingested aggregation protein deposition (pd) [scale bar = 2 μm]. (C) Example of CLEC7A-immunopositive microglial process with extensive ingestion of myelin (im), autophagosomes (x) and a tertiary lysosome (3°) [scale bar = 2 μm]. (D) Magnified image showing the cholesterol crystals (») present in CLEC7A-immunopositive microglia [scale bar = 0.5 μm]. (E) Magnified image of ingested myelin (im) present in a CLEC7A-immunopositive microglia [scale bar = 0.5 μm]. (F) Magnified image showing the protein deposit (pd) aggregation present in a CLEC7A-immunopositive microglia [scale bar = 0.5 μm]. Red outline[=[plasma membrane, blue outline[=[nuclear membrane, orange octothorpe[=[mitochondria, green plus sign[=[endoplasmic reticulum, purple ampersand[= Golgi apparatus, yellow x = autophagosome, yellow » = cholesterol crystals, 3°[=[tertiary lysosome, pd = protein deposition, BV = blood vessel, im = ingested myelin.

Taken together, these observations showed that a variety of disease-associated microglial states displaying ultrastructural markers of cellular stress and extensive interactions with the parenchyma are present in old mice harboring the G2029S pathogenic mutation in Lrrk2.

### CLEC7A Expression is Increased in Primary Lrrk2 G2019S Microglial Cells

To investigate further the expression of CLEC7A in PD pathology, we measured the whole striatal IBA1 and CLEC7A protein contents in Lrrk2 WT and Lrrk2 G2019S mice by Western blot (Suppl. Fig. 4). We found that CLEC7A is heavily detected in the homogenates from both Lrrk2 WT and Lrrk2 G2019S mice, which may suggest that other cells are expressing CLEC7A protein given the low percentage of brain cells represented by microglia (Supplementary Fig. 4). Moreover, the analysis did not reveal significant differences in the overall expression of both IBA1 and CLEC7A in the striata from 18-month-old Lrrk2 WT and G2019S mice (Suppl. Fig. 5A; Lrrk2 WT *vs* Lrrk2 G2019S, all comparisons p > 0.05).

To isolate the contribution of microglial CLEC7A expression, we quantified the amount of the protein in primary microglial cells derived from Lrrk2 WT and Lrrk2 G2019S mice by Western blot. As shown in Fig. 5A, we found that the expression level of IBA1 was comparable between the primary microglial cells derived from Lrrk2 G2019S mice compared to the WT controls (Fig. 5A-B, Lrrk2 WT *vs* Lrrk2 G2019S; p = 0.5915); whereas the expression levels of CLEC7A were enhanced in the primary microglia derived from LRRK2 G2019S mice compared to WT controls (Fig. 5A-C, Lrrk2 WT *vs* Lrrk2 G2019S; p = 0.0022).

**Fig. 5:**
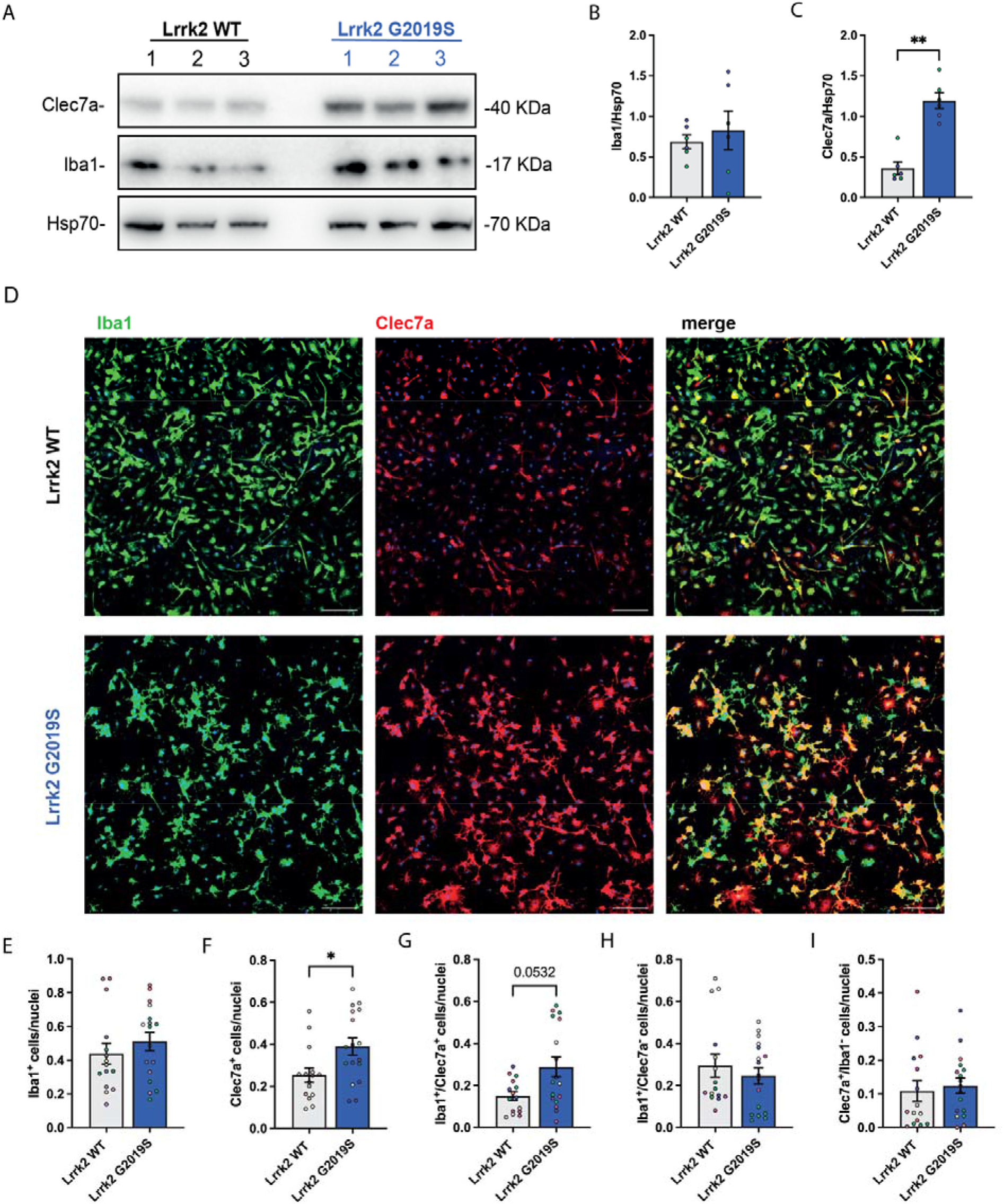
Impact of Lrrk2 G2019S mutation on CLEC7A expression in primary microglial cultures (A) Western blot analysis of primary cortical microglia derived from Lrrk2 WT and Lrrk2 G2019S mice using anti-CLEC7A and anti-IBA1 antibodies; (B-C) Relative quantification of band intensity was performed using ImageJ and normalized to the Hsp70 housekeeping (*n*=6 biological replicates for both Lrrk2 WT and Lrrk2 G2019S); (D) Representative confocal images (20X magnification) of primary cortical microglia derived from Lrrk2 WT and Lrrk2 G2019S mice stained for the microglia and macrophage marker IBA1 (green) and the disease-associated microglial state marker CLEC7A (pseudocolored in red) [scale bar = 100 μm]; (E) Quantification of the number of IBA1^+^ cells, CLEC7A^+^ cells (F), double-positive cells (G), as well as IBA1^+^/CLEC7A^-^ (H) and CLEC7A^+^/ IBA1^-^ (I) cells; data were normalized on the number of counted nuclei. *N* = 5 animals for both Lrrk2 WT and Lrrk2 G2019S were analyzed. Statistical analysis in (F), (G), (I) was performed using an unpaired sample t-test. Statistical analysis in E), H) was performed using a Mann-Whitney U t test.

Additionally, we double stained primary microglial cells with the microglial/macrophage marker IBA1 and CLEC7A (Fig. 5D) and imaged them using confocal microscopy. DM were previously shown to downregulate IBA1, while disease-associated microglial states generally retain this marker (3,7,8). We noticed a slight but not significant increase in the proportion of IBA1^+^ primary microglial cells from Lrrk2 G2019S mice compared to WT controls (Fig. 5D-E; Lrrk2 WT *vs* Lrrk2 G2019S; p = 0.4114). We also quantified a significant increase in the proportion of CLEC7A^+^ cells in Lrrk2 G2019S primary microglial cells compared to the WT counterparts (Fig. 5D, F; Lrrk2 WT *vs* Lrrk2 G2019S; p = 0.0169). Moreover, we did not find any significant differences in the proportion of double positive microglial cells (Fig. 5D, G: Lrrk2 WT *vs* Lrrk2 G2019S; p = 0.0532). Finally, we determined the proportion of IBA1^+^/CLEC7A^-^ (Fig.5D, H) and the proportion of CLEC7A^+^/IBA1^-^ (Fig. 5D, I) microglial cells. In both cases, we did not find significant difference between genotypes (Fig. 5D, H-I: Lrrk2 WT *vs* Lrrk2 G2019S; p = 0.6820; Lrrk2 WT *vs* Lrrk2 G2019S; p = 0.6857). To further investigate the possible relationship between IBA1 and CLEC7A expression, we analyzed the correlation between the integrated density of these two markers. No significant correlation between the CLEC7A and IBA1 integrated density was found (Suppl. Fig. 5B; r < 0.6).

Taken together, these findings show an increase in the proportion of CLEC7A^+^ cells in the *in vitro* primary microglial cell culture derived from Lrrk2 pathogenic mice compared to WT controls.

### CLEC7A-expressing Microglia are more Abundant and Cluster in Aged Lrrk2 G2019S Mice

To further investigate the relevance of CLEC7A^+^ microglia in PD pathology, we next quantified the number of CLEC7A^+^ microglia *in situ* in 18-month-old Lrrk2 WT and Lrrk2 G2019S mice by confocal imaging. To this purpose, we double-stained coronal slices containing the entire DS region (Suppl. Fig. 1A, B) with the markers IBA1 and CLEC7A (Fig. 6A, 6G, 7A). We analyzed the proportion of IBA1^+^-, CLEC7A^+^- as well as IBA1^+^/CLEC7A^+^-microglia (Fig. 6B-D) in this region. We found that IBA1^+^ microglia were densely distributed in the DS of both Lrrk2 WT and Lrrk2 G2019S aged mice, without significant differences between the genotypes (Fig. 6A-B; Lrrk2 WT *vs* Lrrk2 G2019S, p = 0.59). Conversely, we found sparse CLEC7A^+^microglia in the striatum of Lrrk2 WT mice (Fig. 6A, C), while these CLEC7A^+^ cells became abundant in the same region of Lrrk2 G2019S mice (Fig. 6A, C; Lrrk2 WT *vs* Lrrk2 G2019S, p = 0.01), which aligns well with our immunocytochemical ScEM results. Moreover, we quantified the proportion of microglia positive for both markers and we found a significant increase in the proportion of double positive cells (CLEC7A^+^ and IBA1^+^) in the striatum of Lrrk2 G2019S mice compared to WT controls (Fig. 6A, D; Lrrk2 WT *vs* Lrrk2 G2019S, p = 0.004). We also quantified the proportion of IBA1^+^ but CLEC7A^-^, as well as the proportion of CLEC7A^+^ but IBA1^-^ cells *in situ*. In both cases, we did not find significant differences between genotypes (Fig. 6A, E-F; Lrrk2 WT *vs* Lrrk2 G2019S; all comparisons p > 0.05). Moreover, no significant correlation between the integrated density of CLEC7A and IBA1 was found (Suppl. Fig. 5A-B, IBA1 integrated density *vs* CLEC7A integrated density, Pearson’s correlation coefficient R^2^ = 0.02485).

**Fig. 6:**
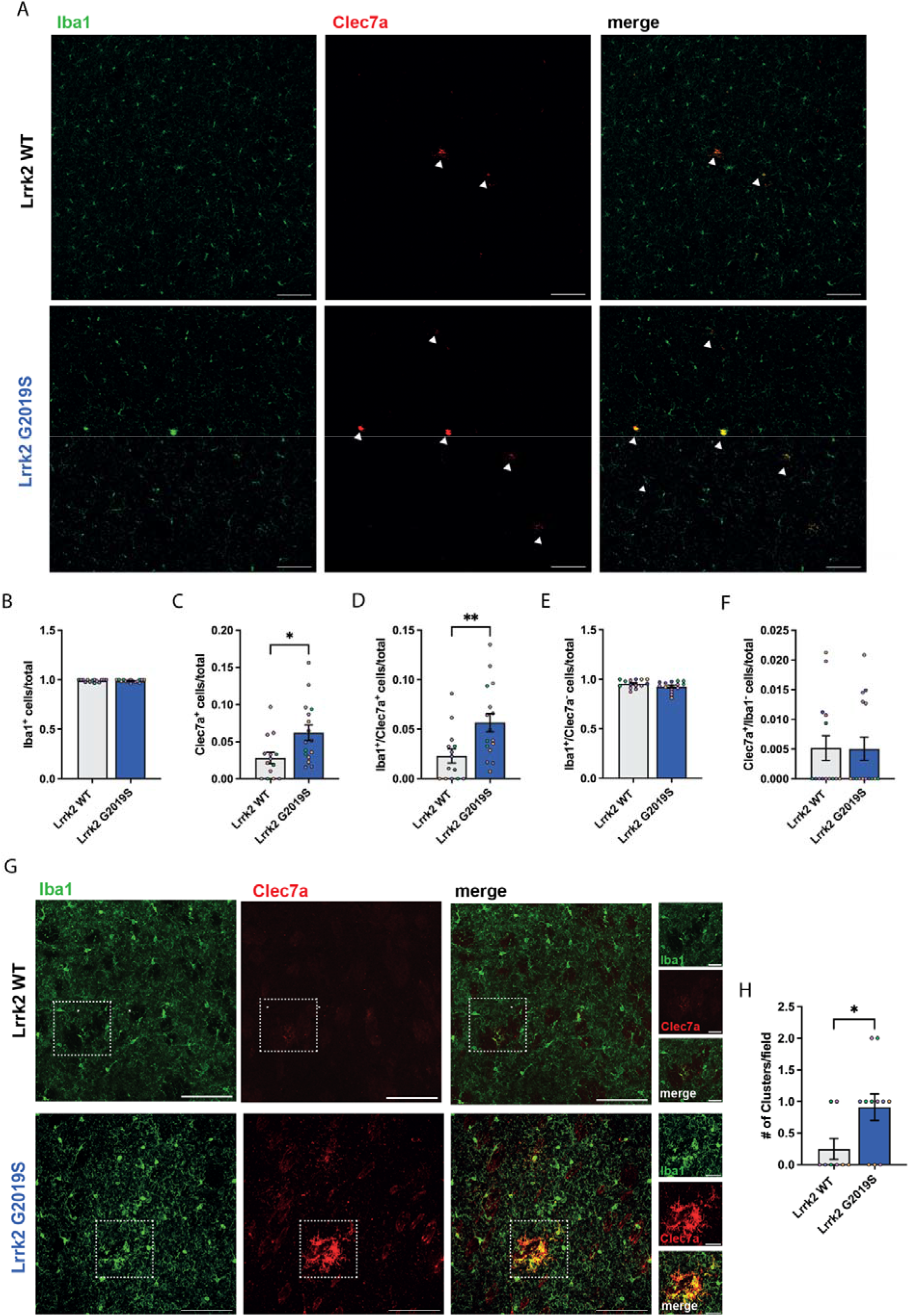
Impact of Lrrk2 G2019S mutation on CLEC7A microglial proportion and clusters in the DS (A) Representative confocal images (20X magnification) of coronal sections derived from 18-month LRRK2 WT and G2019S mice stained for the microglial marker IBA1 (green) and the disease-associated microglial state marker CLEC7A (pseudocolored in red) [scale bar = 100 μm]; (B) quantification of the proportion of IBA1^+^-, CLEC7A^+^-(C), double positive-microglia (D; IBA1^+^+ CLEC7A^+^), as well as IBA1^+^/CLEC7A^-^ (E) and CLEC7A^+^/ IBA1^-^ (F) cells in the DS of 18-months LRRK2 WT and G2019S mice. (G) Representative z-stacks confocal images of CLEC7A-positive clusters (40X magnification) in coronal slices derived from 18-month LRRK2 WT and G2019S mice stained for the microglial marker IBA1 (green) and the disease-associated microglial state marker CLEC7A (pseudocolored in red)-the insets show representative CLEC7A-positive clusters in Lrrk2 WT and G2019S striatal sections [scale bar = 100 μm; insets = 30 μm]; (H) Quantification of the number of CLEC7A^+^ clusters in the striatum of 18-month LRRK2 WT and G2019S mice. The number of clusters were manually counted. *N* = 4 animals for both Lrrk2 WT and Lrrk2 G2019S were analyzed. Statistical analysis in (B), (D), (E), and (H) was performed using a Mann Whitney U test; statistical analysis in (C) and (F) was performed using an unpaired sample t-test.

**Fig. 7:**
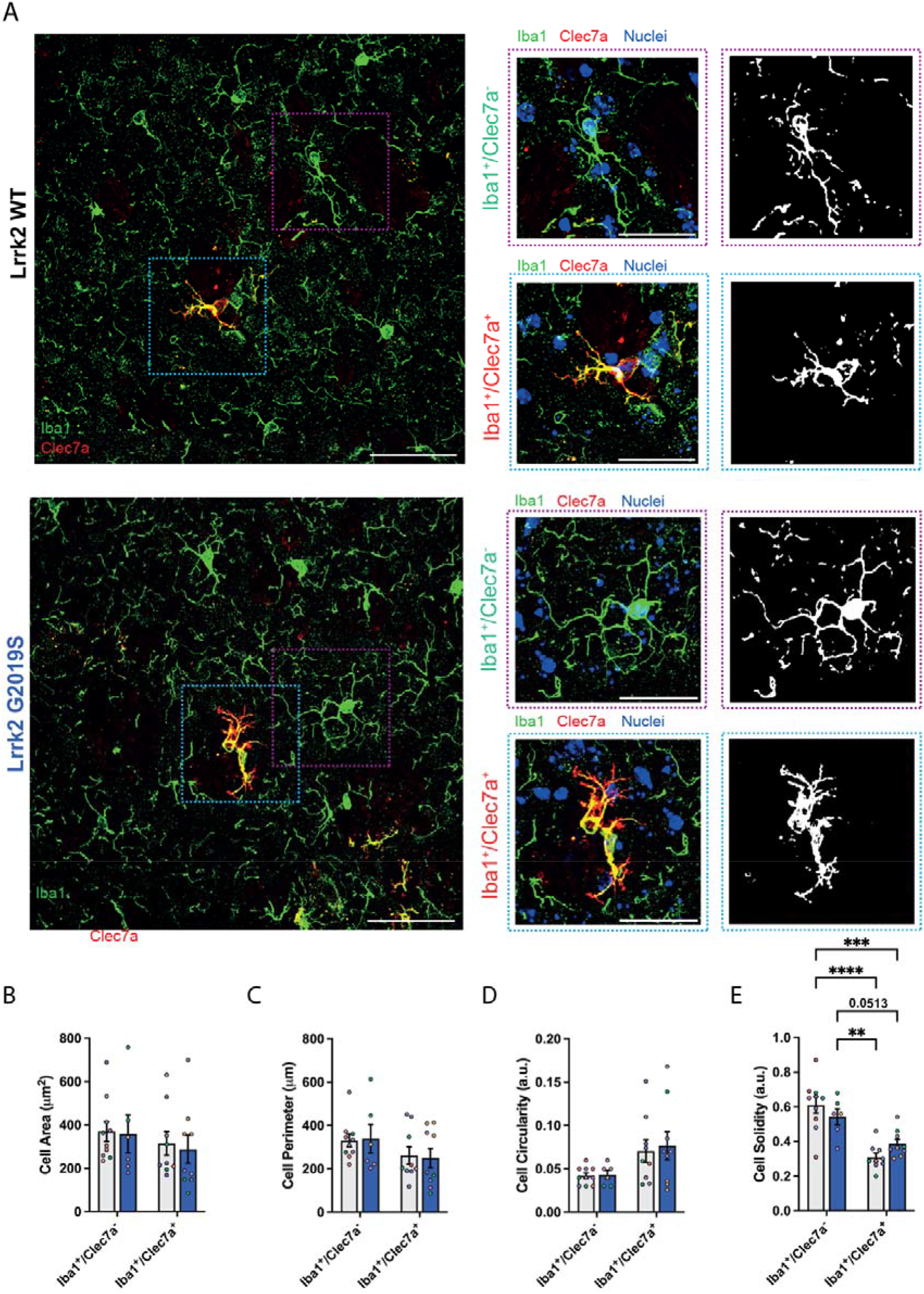
Impact of Lrrk2 G2019S mutation on CLEC7A microglial morphology in the DS (A) Representative z-stacks confocal images (63X magnification) of 18-month-old Lrrk2 WT and Lrrk2 G2019S DS stained for the microglia and macrophage marker IBA1 (green) and the disease-associated microglial states marker CLEC7A (left panels; pseudocolored in red) [scale bar = 50 μm]; insets in purple show IBA1^+^/CLEC7A, while insets in blue indicate representative IBA1^+^/CLEC7A^+^-microglial cells; right panels indicate the relative filtered binary mask [scale bar = 30 μm]; (B) Quantification of the measured cellular area, the cellular perimeter (C), the circularity (D) and the solidity index of the analyzed microglia. (B-E) *n* = 4 mice for each genotype. Statistical analysis in (B-E) was performed using a two-way ANOVA test followed by Tukey’s multiple comparisons test.

To provide additional insights into the CLEC7A^+^ microglial state, we imaged at higher magnification the CLEC7A^+^ cells (IBA1^+^ and IBA1^-^), and we noticed that CLEC7A^+^ displayed a pronounced clustered phenotype in the Lrrk2 G2019S mice (Fig. 6G). By quantifying the number of CLEC7A^+^ clusters in the DS, we found no (or sporadic) CLEC7A^+^ clusters in the striatum of Lrrk2 WT mice (Fig. 6G-H); conversely, we found a significant increase of clustering in the Lrrk2 G2019S pathological model (Fig. 3H; Lrrk2 WT *vs* Lrrk2 G2019S, p = 0.04). Similarly, the DM were previously shown to form clusters (3), a finding that was reproduced in the current study, in the context of PD pathology (Fig. 3C).

Taken together, these findings indicate a genotype-dependent increase in the proportion of CLEC7A^+^ microglia which tend to amass in presence of the Lrrk2 pathological mutation.

### CLEC7A-positive Microglia Display an Ameboid Morphology in Aged Lrrk2 G2019S Mice

To dissect the impact of CLEC7A expression on microglial structure and function, we further examined possible differences in morphology between IBA1^+^/CLEC7A^-^ and IBA1^+^/CLEC7A^+^ cells in the DS of aged Lrrk2 G2019S mice *versus* controls (Fig. 7; A-E). We measured different morphological parameters of the entire process arborization: the cell area, perimeter, circularity and solidity (Fig. 7 B-E). We found no changes in the area, perimeter and circularity between the IBA1^+^/CLEC7A^-^ and IBA1^+^/CLEC7A^+^ cells and among the genotypes (Fig. 7 B-D; Lrrk2 WT *vs* Lrrk2 G2019S all comparisons p > 0.05). However, we found a significant genotype independent decrease in solidity between IBA1^+^/CLEC7A^+^ cells and IBA1^+^/CLEC7A^-^ cells (Fig. 7A, E, IBA1^+^CLEC7A^-^: Lrrk2 WT *vs* IBA1^+^CLEC7A^+^: Lrrk2 WT, p < 0.0001; IBA1^+^CLEC7A^-^: Lrrk2 WT *vs* IBA1^+^CLEC7A^+^: Lrrk2 G2019S, p = 0.0005; IBA1^+^CLEC7A^-^: Lrrk2 G2019S *vs* IBA1^+^CLEC7A^+^: Lrrk2 WT, p = 0.0017; IBA1^+^CLEC7A^-^: Lrrk2 G2019S *vs* IBA1^+^CLEC7A^+^: Lrrk2 G2019S, p = 0.0513). Finally, the Sholl analysis performed on the IBA1^+^/CLEC7A^-^ and IBA1^+^/CLEC7A^+^ cells suggest a higher morphological complexity of the IBA1^+^/CLEC7A^-^ population in the proximal region compared to the IBA1^+^/CLEC7A^+^ cells (Supp. Fig 5E, IBA1^+^CLEC7A^-^ *vs* IBA1^+^/CLEC7A^+^, p = 0.0005). These data further suggest that CLEC7A expression in microglia promotes a shift away from their homeostatic state.

## Discussion

In this work, we investigated for the first time the diversity of microglial states in aged mice harboring a G2019S mutation in the PD-linked LRRK2 gene. We focused our cellular and ultrastructural characterization on the striatum, one of the brain regions of special relevance to PD accordingly to the dying back hypothesis. Indeed, Tagliaferro and Burke work suggests that neurodegeneration of the dopaminergic neurons might start from the neuronal terminals located in the striatum rather than the cell bodies in the SNpc (54). Here, we identified several ultrastructural markers of pathology in striatal microglia as well as the presence of two disease-related microglial states, the dark and CLEC7A^+^ disease-associated microglia. Additionally, we developed and employed a novel integrated density approach to measure microglial electron density through ScEM, which allowed us to distinguish between dark, intermediate, and typical microglia quantitatively. This approach provides categorical consistency and control for subtle differences in individual electron micrograph brightness and contrast properties. It can be applied to further research investigating ‘dark cell’ populations.

Using extensive ScEM analyses, we demonstrate a significant impact of the LRRK2 G2019S mutation on the ultrastructural characteristics of microglia and their interactions with the neuropil in the striatum. Significant increases in ER and Golgi body dilation were observed in microglia from animals expressing LRRK2 mutation. Dilation in these organelles is known to indicate stress via the accumulation of misfolded or unfolded proteins (54,55), suggesting a microglial population-wide alteration in organellar health associated with the LRRK2 G2019S mutation. Similar ultrastructural alterations in microglial ER and Golgi stress have been described in a mouse model of Alzheimer’s disease pathology (4). Furthermore, identifiable contacts of the LRRK2 G2019S microglia with the surrounding neural tissue were significantly decreased compared to WT. Moreover, we observed a significant decrease in the number of contacts between microglia and the axon terminals but not in interactions with post-synaptic dendrites or neuronal cell bodies. Our observations indicate that this is likely due to an increased phagocytic activity as LRRK2 G2019S microglia were contacting significantly more extracellular digestion and space. Phagocytosis of neural debris and clearance of damaged organelles are essential physiological functions of microglia which are critical to healthy neurodevelopment and maintaining brain health throughout the lifespan (56–58). However, in the contexts of chronic stress, infection, aging, and degeneration, for example, microglia may begin to over-phagocytose the parenchyma resulting in behavioral, cognitive, and emotional deficits (59–63).

Additionally, the current study identified a significantly increased presence of darker microglia and presented the first visualization of CLEC7A^+^ microglia in the context of PD pathology, in the striatum of LRRK2 G2019S mice. CLEC7A has been identified as a marker for disease-associated microglial states (7,8). Upon visualization of these cells through electron microscopy, we observed substantial evidence that these CLEC7A^+^ microglia undertake severe phagocytic activity, particularly by the presence of extensive phagosomes, and accumulate protein aggregates which is slightly enhanced in the striatum of these mice (64). The CLEC7A^+^ microglia had both typical and intermediate nuclear patterns and did not meet the classic criteria for DM—suggesting that DM may downregulate CLEC7A in the current context, although the immunostaining might have masked the cell’s electron density, which warrants further examination. As the burden on these disease-associated cells exceeds capacity, this may lead to a phenotypic change and expression of the canonical features of DM (nuclear heterochromatin loss, ultrastructural stress markers, and mitochondrial cristae alterations). Altogether, we demonstrate that substantial neural degeneration as a result of the G2019S mutation on the LRRK2 gene leads to a highly-stressed microglial state in the aged brain of mice—evidenced by ultrastructural features of cellular stress, increased contacts with extracellular degeneration, and an accumulation of active phagosomes and protein accumulation observed in CLEC7A^+^ microglia—likely contributing to the neurodegenerative cascade of PD.

In IF assays, we did not detect any changes in the expression of the microglial marker IBA1 in the whole brain of the aged mice as well as in the primary microglial cells from Lrrk2 G2019S mice compared to Lrrk2 WT control. On the contrary, CLEC7A protein was abundantly and equally expressed in the striatum of pathogenic and control mice, but it drastically increased in cultured microglial cells from Lrrk2 G2019S mice *versus* Lrrk2 WT control. Overall, these observations suggest that CLEC7A expression might be pathologically relevant only in microglia. Focusing on the IBA1^+^/CLEC7A^+^ cells, we found that they were sparse but enhanced in the DS of the Lrrk2 G2019S mice, compared to the Lrrk2 WT control. This is in line with prior studies that identified CLEC7A-expressing microglial states in other neurodegenerative conditions, such as AD, ALS, and MS (7,8,65). In addition, the presence of CLEC7A-expressing microglia in the WT mice could be due to aging. This hypothesis is supported by previous transcriptome studies that found microglial clusters overexpressing CLEC7A in aged mice (9–11). To confirm the hypothesis of an age-dependent increase in striatal CLEC7A^+^ microglia, we aim to characterize in the future this microglial state also in younger Lrrk2 G2019S and Lrrk2 WT control mice.

We further revealed that the CLEC7A^+^ (IBA1^+^ and IBA1-) microglia form clusters in the DS of the 18-month-old mice. Our findings are in agreement with previously published data that described the clustering of CLEC7A^+^ in a mouse model of another neurodegenerative condition, AD pathology (8,66,67). Similarly, DM were found to form clusters in the current study (Fig. 3C) and previous work in a mouse model of AD pathology (3). As a future prospective, the CLEC7A-associated microglial state can be investigated also in a murine model that recapitulates the late stage of PD or in the Lrrk2 G2019S model treated with compounds able to trigger neurodegeneration (i.e., *in vivo* alpha-synuclein injection). Microglial shift from homeostasis is usually associated with a series of concerted morphological and functional changes (68,69). To provide insights into the roles of CLEC7A microglia, we assessed multiple morphological parameters of CLEC7A-expressing cells compared to homeostatic IBA1^+^/CLEC7A^-^ microglia. We found that the cell solidity was significatively different between the two microglial states. The solidity index describes the stiffness and deformability of an object (50). In particular, the higher the solidity, the lower the cell deformability. Lower solidity values in the CLEC7A microglia indicates that these cells are highly deformable, which in turn suggests an increase in their motility and migration. In fact, as previously described, in the presence of dangerous stimuli, microglia generally increase their migration toward the site of inflammation (61). In accordance, Moehle and colleagues demonstrated that the Lrrk2 G2019S peripheral myeloid cells exhibit increased motility, whereas the pharmacological inhibition of the kinase activity of Lrrk2 blocked the enhanced chemotaxis associated with the G2019S mutation (20). Moreover, the Lrrk2 G2019S fibroblasts have an enhanced motility compared to Lrrk2 WT control (70). Consistently, the CLEC7A microglia might migrate and surround the source of inflammation, such as extracellular debris requiring removal. Indeed, our ultrastructural analysis of the CLEC7A microglia indicate a high presence of phagosomes containing myelin and we confirmed by immunocytochemistry and Western blot the expression of CLEC7A in the primary culture.

In the striatum of the 18-month-old mice, we found 4–6% of CLEC7A positive cells, whereas in the primary microglia we found 20–40% of CLEC7A microglia. This result might be explained by the fact that the primary microglial cultures were obtained from pups. Indeed, scRNAseq analysis has shown that CLEC7A is overexpressed in the proliferative-region associated microglia (PAM) acutely isolated from mice at postnatal day 7 (71). This data suggests that CLEC7A might be highly present in microglial cells at the early stages of life. Moreover, the primary microglial cultures used in our experiments derive from the cerebral cortex, a region where CLEC7A^+^ cells could be overrepresented. Specifically, scRNAseq data indicates that acutely isolated cortical microglial cells called white matter-associated microglia (WAMs) upregulate genes involved in phagocytic activity and lipid metabolism, including CLEC7A (72).

## Conclusions

Here, we used various imaging and analysis techniques to investigate microglial states in the context of a mouse model with a G2019S mutation of the LRRK2 gene. LRRK2 G2019S mutation is the most common mutation associated with sporadic PD in humans. Ultrastructural investigation of microglia in the DS demonstrated population-wide cellular stress as well as increased microglia-associated extracellular digestion and space pockets. Striatal CLEC7A^+^ microglia were visualized using ScEM, showing that they displayed extensive phagosomes but were not overlapping with DM which were also significantly more abundant in G2019S mutation mice. Further, the CLEC7A^+^/IBA1^+^ cells were morphological different compared to CLEC7A^-^/IBA1^+^ cells. CLEC7A+ cells tend to form clusters *in situ*, while IBA1^+^ cells overexpressed CLEC7A in primary microglial cultures from LRRK2 G2019S mice. Altogether, these data indicate that multiple disease-linked microglial states coexist in the Lrrk2 G2019S mouse striatum which may contribute together to PD pathology.

## List of Abbreviations

AD: Alzheimer’s disease

ALS: amyotrophic lateral sclerosis

CNS: central nervous system

DAM: disease associated microglia

DM: dark microglia

DMEM: Dulbecco’s Modified Eagle Medium

DS: dorsal striatum

DSp: periventricular dorsal striatum

ER: endoplasmic reticulum

FBS: fetal bovine serum

FIB-SEM: focused-ion beam scanning electron microscope

HD: Huntington’s disease

HRP: horseradish-peroxidase

IF: immunofluorescence

IHC: immunohistochemistry

Il-: Interleukin-

MGnD: microglial neurodegenerative phenotype

MS: multiple sclerosis

PAM: proliferative-region associated microglia

PB: phosphate buffer

PBS: phosphate-buffered saline

PD: Parkinson’s disease

PFA: paraformaldehyde

ROS: reactive oxygen species

RT: room temperature

ScEM: scanning electron microscopy

SEM: standard error of the mean

SNpc: substantia nigra pars compacta

TBS: tris-buffered saline

TBS-T: Tris-buffered saline containing 0.1% Tween

TNF-α: tumor necrosis factor-alpha

WAM: white matter-associated microglia

WT: wild-type

## Declarations

### Ethics approval and consent to participate

Housing and handling of mice were done in compliance with national guidelines. All procedures performed with mice were approved by the Ethical Committee of the University of Padova and the Italian Ministry of Health (license 200/2019).

### Consent for publication

Not applicable

### Availability of data and materials

The datasets used and/or analyzed during the current study are available from the corresponding author on reasonable request.

## Authors’ contributions

MET and LC designed the study, interpreted data, and wrote the manuscript; LI, JV, KG and KP performed and optimized the experiments, interpreted data, and wrote the manuscript; VG and AC helped with the preparation of the samples; LI and KG performed immunofluorescence and Western blot experiments; JV and MK processed tissue for EM, imaged on the FIB-SEM, and analyzed EM data; LI, JV and GK prepared figures; EG has contributed to manuscript editing and optimization. LTL contributed to arranging shipment and sample processing. All authors have read and approved the final manuscript.

## Acknowledgments

We acknowledge and respect that the University of Victoria is located on the territory of the lək[wəŋən peoples and that the Songhees, Esquimalt, and WSÁNEÆ peoples have relationships to this land. University of Padova to support LC as assistant professor and the IRCCS San Camillo Hospital in Venice, Italy.

## Authors’ Information

LI, JV, and GK share the first co-authorship. MET and LC are co-corresponding authors.

---

**Supp. Fig. 1:**
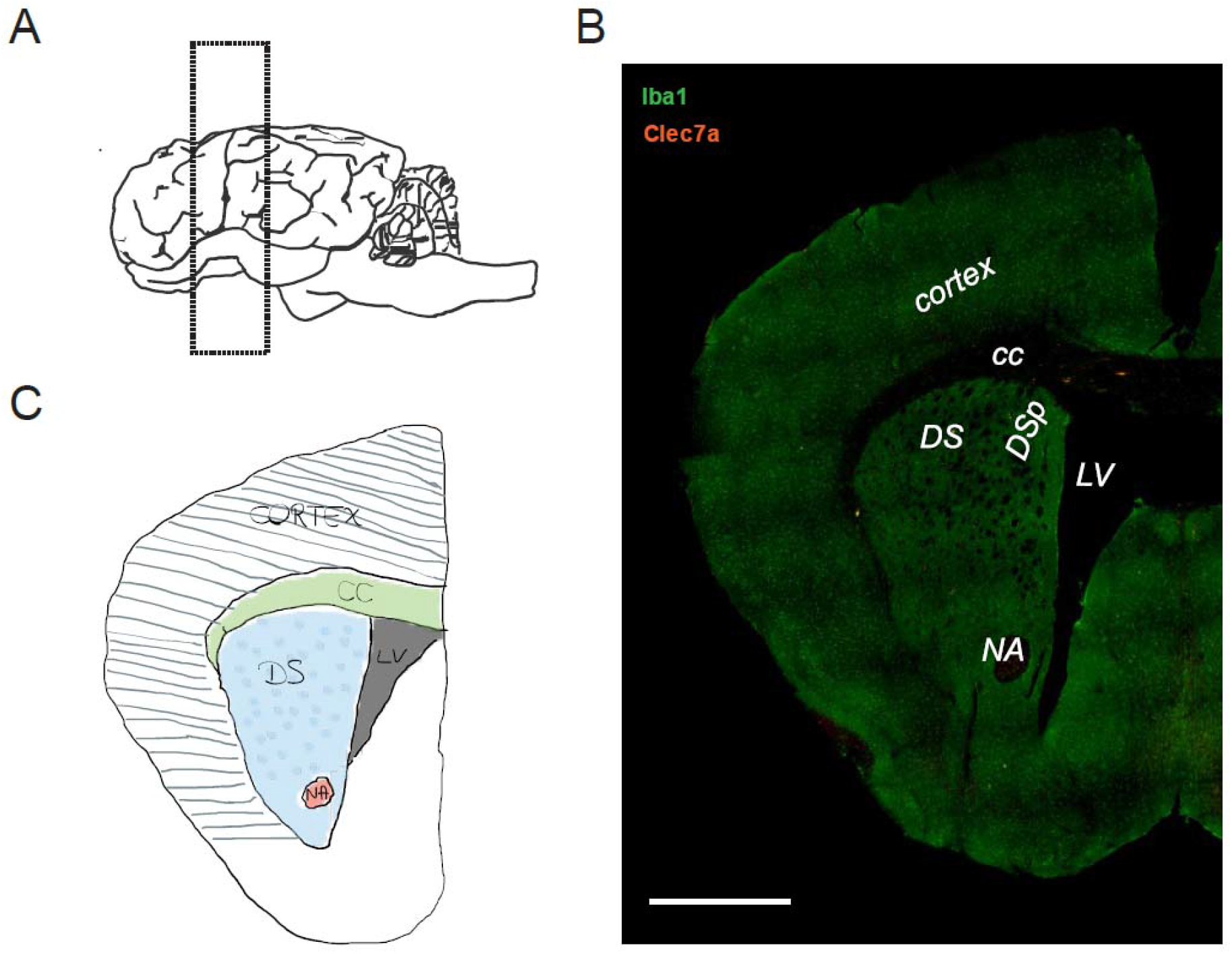
Anatomical region of interest displayed by a representative cartoon and an immunofluorescence overview (A) Representative cartoon of the dorsal striatum (DS) region considered in our analysis and of the coronal slice anatomy [scale bar = 1mm] (B). (C) Representative epifluorescence image of a coronal slice containing the DS. In the section, the DS can be identified by the presence of several anatomical hallmarks such as its location immediately ventral to the corpus callosum (CC) and to the cortex, dorsal to the nucleus accumbens (NS) and close to the lateral ventricle (LV). The DS region immediately lateral to the LV is indicated as the dorsal striatum periventricular (DSp).

**Supp. Fig. 2:**
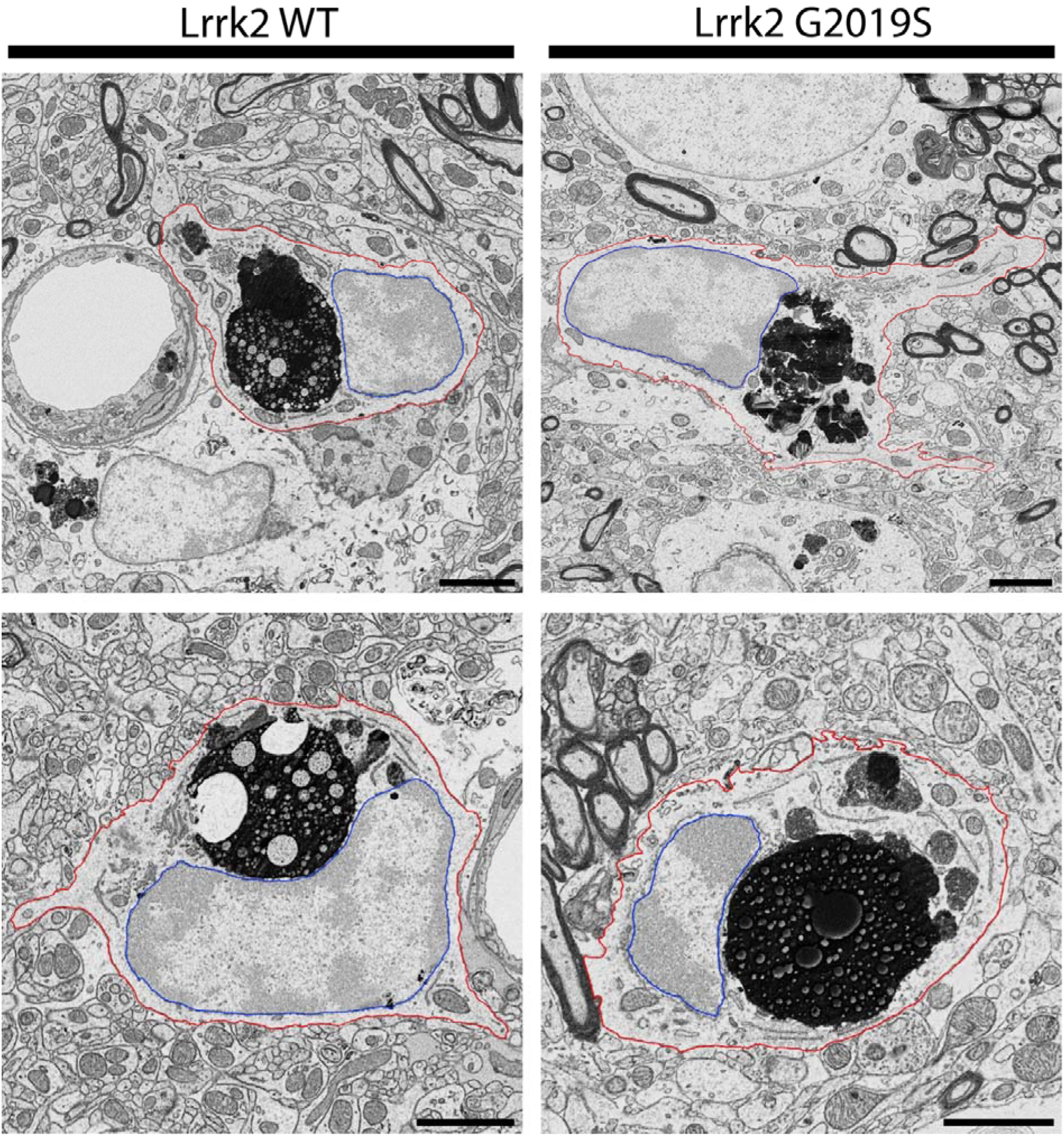
Exemplar electron micrographs showing microglia in the periventricular dorsal striatum of Lrrk2 WT and G2019S animals. These exemplar microglial cell bodies are imaged at 5 nm-resolution in x-and-y and are representative of the other images used in ultrastructural analysis [scale bar = 2 μm]. Red outline[=[plasma membrane, blue outline[=[nuclear membrane.

**Supp. Fig. 3:**
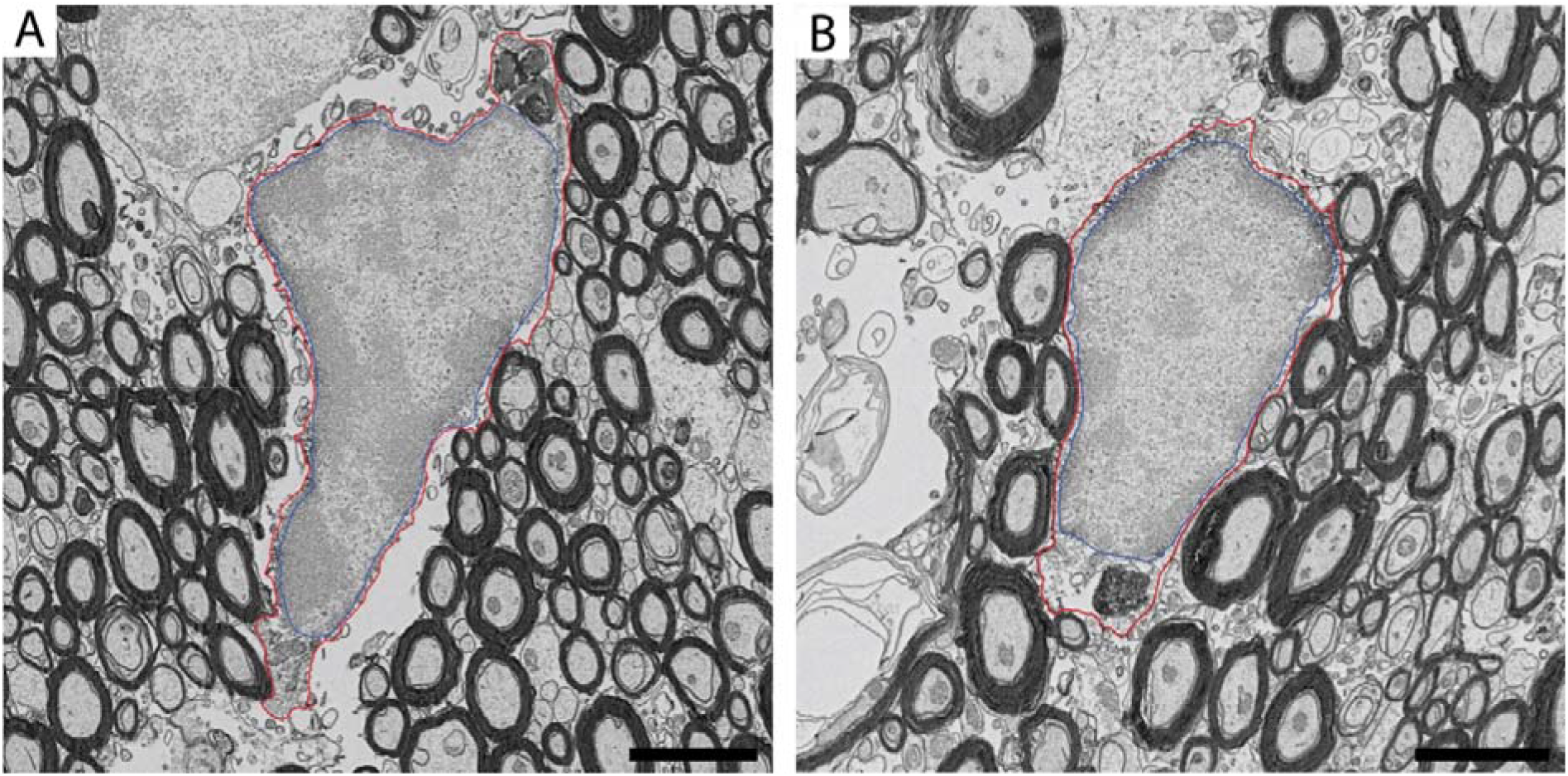
Examples of CLEC7A-positive microglia displaying intermediate nuclear patterns. (A-B) Examples of CLEC7A-immunopositive microglial cell bodies which display intermediate nuclear patterns and nucleoplasmic electron density, suggestive of intermediate microglia, in the DSp of a Lrrk2 G2019S mouse [scale bar = 2 μm]. Red outline[=[plasma membrane, blue outline[=[nuclear membrane.

**Supp. Fig. 4:**
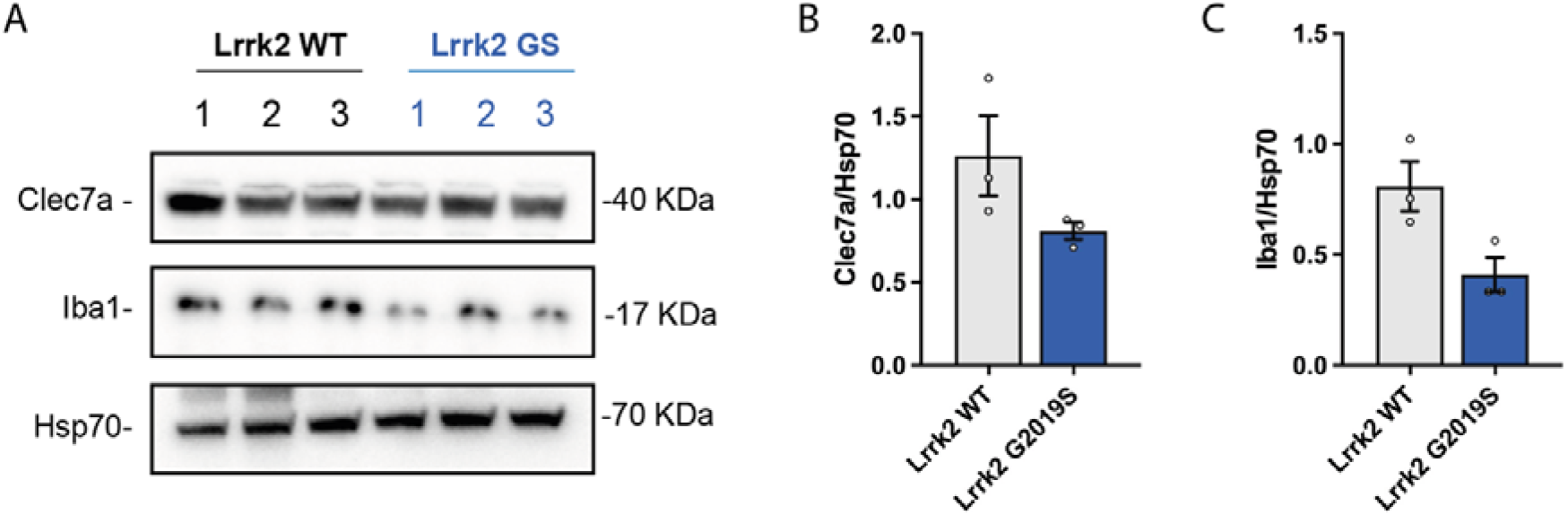
CLEC7A expression in the striatum lysate of 18-month-old Lrrk2 WT and Lrrk G2019S mice (A) Western blot analysis of LRRK2 WT and LRRK2 G2019S striatal lysates using anti-IBA1 and anti-CLEC7A antibodies; (B, C) Relative quantification of band intensity was performed using ImageJ and normalized to the Hsp70 housekeeping; *n* = 3 striatal samples for both Lrrk2 WT and Lrrk2 G2019S 18-month-old mice. Statistical analysis in (B) was performed using an unpaired sample t-test while in (C) was performed using a Mann Whitney U test.

**Supp. Fig. 5:**
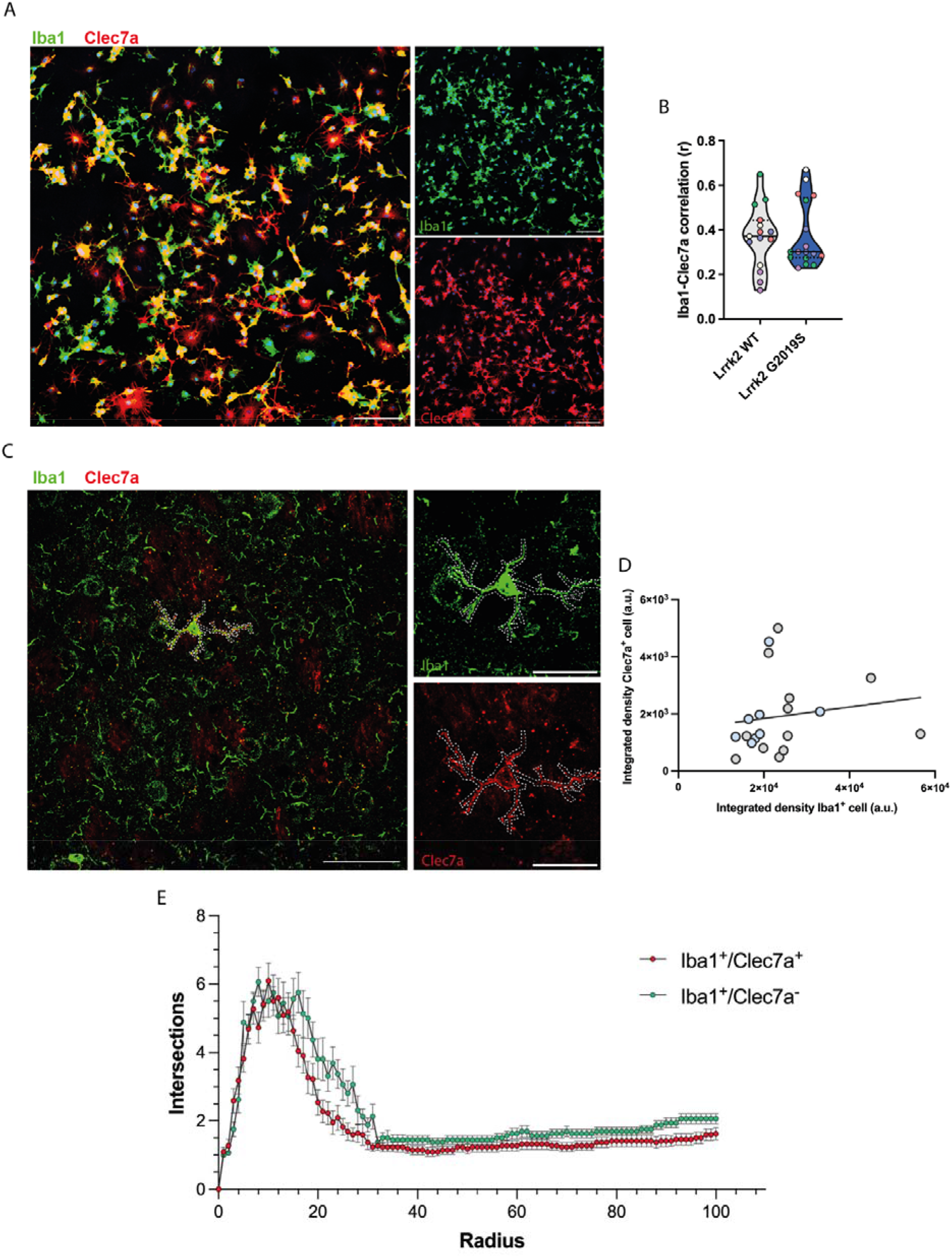
Integrated density of CLEC7A and IBA1 *in vivo* and *in vitro* (A) Representative confocal image (20X) showing primary microglial cells derived from Lrrk2 WT and Lrrk2 G2019S mice [scale bar = 100 μm] (B) Correlation analysis between IBA1 and Dectin1 was performed in both genotypes; *n*=5 microglial cells derived from 5 independent cultures. (C) Representative confocal z-stack image (63X magnification) in the dorsal striatum of 18-month-old mice with the region of interest around IBA1^+^-CLEC7A^+^ cells [scale bar = 100 μm; scale bar insets = 30 μm]. Integrated density of the ROI for both IBA1 and CLEC7A was evaluated; n = 4 per genotype (D) Quantification of the integrated density of IBA1 and CLEC7A in microglia localized in the dorsal striatum of both Lrrk2 WT mice (grey points) and Lrrk2 G2019S mice (blue points). Statistical analysis in (B) was performed using an unpaired sample t-test to compare the correlation between the two genotypes. In (D) a linear regression analysis was performed. (E) Results of sholl analysis performed on IBA1^+^/CLEC7A^+^ and IBA1^+^/CLEC7A^-^ cells (step size = 1 μm; radius = 100 μm). Supplementary Figure 5: Representative microglial micrographs from G2019S and WT mice.

## Supplementary

Supp. Table 1: Quantitative Ultrastructural Analysis of Microglia Results Table *Attached as an excel sheet (as per journal guidelines)*

